# Piercing mechanism in the minimal contractile tail of bacteriophage P2

**DOI:** 10.64898/2026.09.14.750843

**Authors:** Victor Klein-Sousa, Marta Šiborová, Aritz Roa-Eguiara, Monica Santiveri, Inga Songailiene, Nicholas M. I. Taylor

**Affiliations:** Novo Nordisk Foundation Center for Protein Research, Department of Cellular and Molecular Medicine, Faculty of Health and Medical Sciences, University of Copenhagen, Blegdamsvej 3B, 2200 Copenhagen, Denmark; Department of Pharmacology and Drug Design, Faculty of Health and Medical Sciences, University of Copenhagen, Universitetsparken 2, 2100 Copenhagen, Denmark; Institute of Biotechnology, Life Sciences Center, Vilnius University, Vilnius, Lithuania

**Keywords:** Phage P2, Contractile Injection Systems, Cryo-EM, Cryo-ET, Structural Virology

## Abstract

Bacteriophage P2 is a long-standing model for contractile injection systems, yet high-resolution structures of the intact virion and the mechanism of baseplate activation have remained unresolved. Here, we combine high-resolution cryo-electron microscopy and *in situ* cryo-electron tomography to characterize P2, using its lytic mutant P2*vir*, in its pre- and post-contraction states and during host infection. We identify a previously undescribed fold in the head-to-tail adaptor, which mediates the transition from 12-fold portal symmetry to sixfold tail symmetry through interlocking β-strand “handshake” interactions. Our structures further reveal a compact, threefold-symmetric baseplate established by a distinctive loop-helix module formed by the hub and tube initiator, which bioinformatic and phylogenetic analyses identify as a characteristic architectural feature of the *Peduovirus* lineage. We also propose a rope-like arrangement that anchors the tail fibre to the baseplate triplex component gpI and obtain evidence that the tape-measure protein adopts a C3 arrangement within the tail lumen, while computational predictions suggest a possible hexameric, pore-like organization after release. Comparison of pre- and post-contraction particles, supported by *in situ* imaging of infection, suggests that sheath contraction destabilizes the baseplate, promoting its disassembly. The inner tail tube appears to penetrate the host outer membrane but does not appear to reach the inner membrane, leading us to propose that the reorganized tape-measure protein forms a secondary conduit for genome translocation. These findings support a distinct model of contractile tail activation and expand the known architectural diversity of viral injection systems.

## Introduction

Bacteriophages (or phages) are engaged in a constant evolutionary arms race with bacteria, which has equipped phages with specialized mechanisms to hijack the host replication machinery, often leading to cell death^1–3^. As a result, phages are increasingly considered promising antimicrobial agents^4–6^. Phage-host specificity is largely determined by mechanisms of host recognition and genome delivery, both of which depend on the architecture of the phage virion^7–10^.

Myoviruses are long-tailed phages that employ a contractile mechanism to breach bacterial membranes^11^. The structures of distantly related myoviruses have previously been solved by cryo-electron microscopy (cryo-EM), spanning a wide range of genome sizes^12–17^. Large-genome myoviruses include phages P1, φTE, T4, and φ812, whereas small-genome myoviruses include phages such as Mu and Pam3, the latter encoding only 17 genes that form the virion components. As it is common to all these phages, the virion comprises an icosahedral capsid, which is formed by the major capsid protein. The capsid is connected to the tail through a large macromolecular assembly, often referred to as the neck. The tail trunk consists of two layers: an inner tube, which frequently displays density within its lumen, generally attributed to the tape-measure protein (TMP)^13,17,18^, and an outer helical sheath. The sheath is assembled in an extended, high-energy state and contracts to drive penetration of the bacterial outer membrane^19^. At the distal end of the tail, the baseplate forms another macromolecular assembly that contains proteins responsible for host recognition, including receptor-binding proteins, typically organized as fibre-like structures, as well as proteins that trigger sheath contraction and peptidoglycan-cleaving enzymes^14,20,21^. Despite this conserved architecture, mechanisms for host-recognition and contraction can be highly distinct among different phage families ^21^. *Peduovirus* P2 is a well-characterized myovirus^22^, that has large biotechnological and biomedical applications^23–25^. P2 is an *Escherichia coli* phage, first isolated in 1951 by Giuseppe

Bertani^26^. P2 can infect *E. coli* K12 and C strains, and exhibits temperate behaviour, capable of entering either a lytic or lysogenic cycle^22,27^. P2 is the most extensively studied member of the *Peduoviridae* family, which currently comprises 58 genera^28^. All members of this family share a set of six orthologous genes, encoding two enzymatic proteins, the large and small subunits of terminase, and four structural proteins, the portal protein (gpQ), capsid scaffolding protein, major capsid protein (gpN), and capsid completion protein (gpL). These genes are integral to DNA packaging and capsid assembly, respectively. Despite this conservation, *Peduoviridae* genomes vary in size from approximately 28 kb to 41 kb, with P2 possessing a 33,592 bp genome containing 42 open reading frames^22,29^.

The structure of P2’s tail has been first described by transmission electron microscopy in the 70s^30^, which identified six key proteins involved in tail maturation, including the tail fibre protein gpH and the tape-measure protein gpT. Subsequent studies proposed additional components of the baseplate and tail^31,32^, as well as the dodecameric portal protein gpQ^33^, and identified gpU as a putative structural protein whose location within the virion remained unknown^22^.

Previous studies have identified parallels between P2 and R-type tailocins^34–36^, bacterially produced toxin systems that resemble the contractile tails of myoviruses. High-resolution structures of pyocin R2 hint at homology with P2 tail sheath, inner tube, and several baseplate proteins, supporting a shared evolutionary origin^37^. However, the two systems differ in baseplate organization: pyocin R2 relies on a ripcord domain within the tube initiator protein to stabilize the pre-contracted state and mediate contraction, whereas P2 lacks a ripcord domain, implying a distinct mechanism of baseplate assembly and activation.

In this study, we use cryo-EM to determine high-resolution structures of bacteriophage P2*vir*, a strictly lytic mutant of phage P2. We resolve key components of the tail, portal, baseplate, and tail fibre in both pre- and post-contraction states, providing structural insight into tail diversity within *Peduoviridae* phages and contractile injection systems. Moreover, we applied cryo-electron tomography (cryo-ET) to resolve the interaction of P2 with its host. We propose a contraction mechanism that relies on the breaking of the baseplate to control the piercing distance which allows transduction of viral DNA.

## Results and discussions

### An overview of P2 global structure

We resolved the pre- and post-contraction virion structural states from the cryo-EM dataset via local reconstruction of the phage subregions. Within our cryo-EM datasets, we found 2D-classes for both pre- and post-contraction states (Supplementary Fig. 1-3). We processed each state independently. The virion architecture of bacteriophage P2 can be divided into five major structural regions: the capsid, neck, tail trunk, baseplate, and tail fibres (Fig. 1). We reconstructed each of these virion components separately with the appropriate symmetry imposed, and atomic models were built for the asymmetric units, in both pre- and post-contraction states where possible (Supplementary Fig. 1-3)

**Figure 1.**
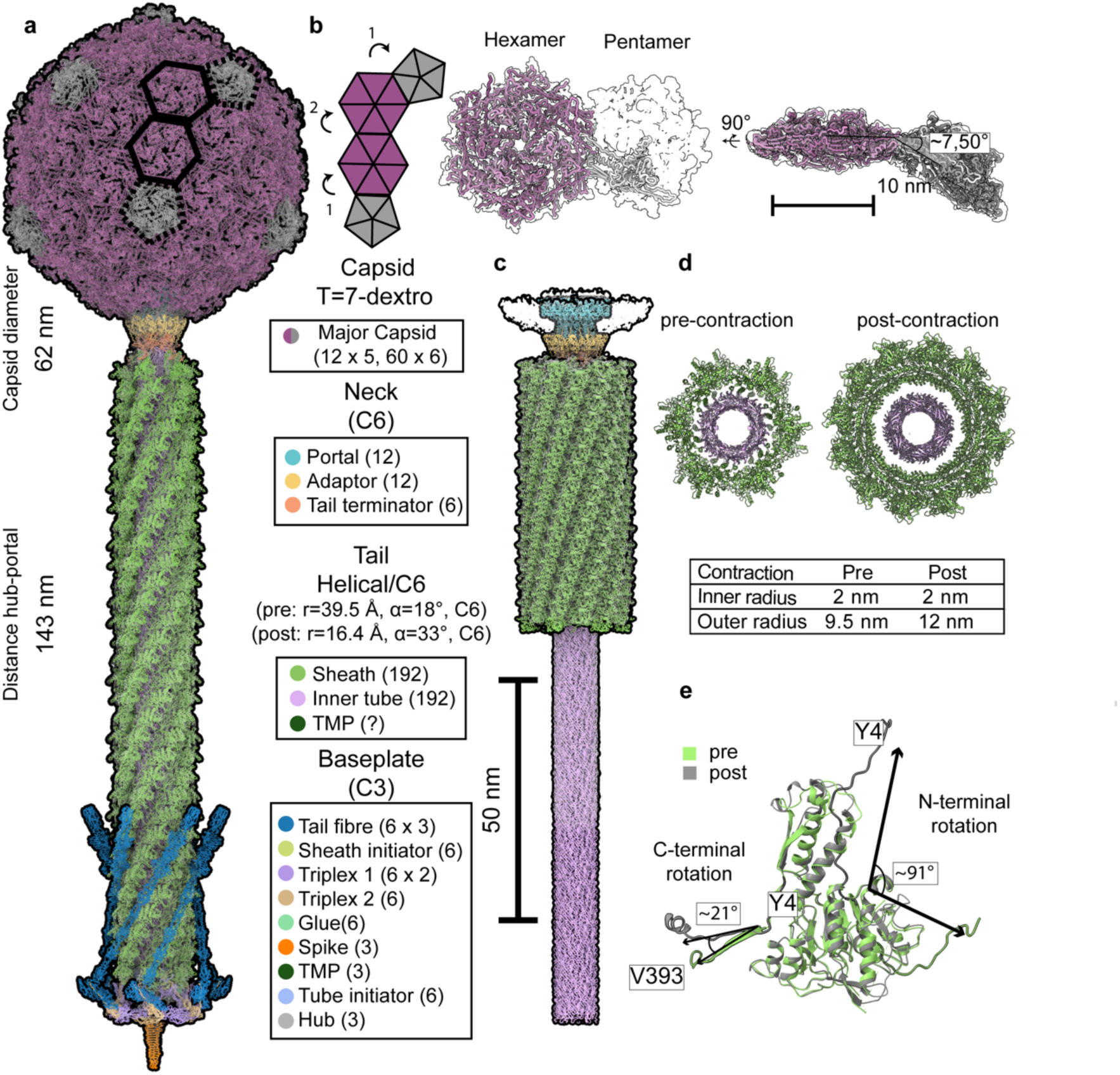
Structure of *Peduovirus* P2 in its pre and post contraction states. (a) Composite model generated from locally resolved maps (capsid: PBD ID 29NY / EMD-57258, contour level 0.13; neck: PBD ID 29NE / EMD-57255; baseplate: PBD ID 29ND / EMD-57254; fibres: EMD-57256). (b) Capsomer map and model. (c) Composite model of the post-contraction virion derived from locally resolved maps (neck and apical tail: EMD-57344; basal tail and inner tube EMD-57343). (d) Horizontal cross-section of the pre- and post-contraction tail showing the diameters of the inner tube and sheath. (e) Model for sheath protein gpFi showing terminal rotations pre-/post-contraction.

The P2 capsid is formed by the major capsid protein gpN and adopts a T=7 dextro icosahedral symmetry, consistent with previous reports^38^ (Fig. 1b). No clear density was resolved for the scaffolding protein O* in the icosahedral map, which agrees with earlier studies indicating that O* is likely disordered in the head and only interacts transiently with gpN during assembly^38–40^. The major capsid protein follows a similar fold as those described for the capsid proteins of other T=7 capsids, such as from phage T7^41^. This consists in an N-arm, E-loop and A-domains (Supplementary Fig. 3). The asymmetric vertex of the capsid binds to the neck to join the sheath and tube (Supplementary Fig. 4). The neck is composed of three rings. The first ring consists of 12 copies of the portal protein Q*^39^, the N-terminally cleaved version of gpQ, for which we could construct an atomic model for residues 27-340. The second ring is made of 12 copies of the head-to-tail adaptor protein gpL, and the final ring consists of six copies of the tail terminator protein gpR (Fig. 1, Fig 2).

**Figure 2.**
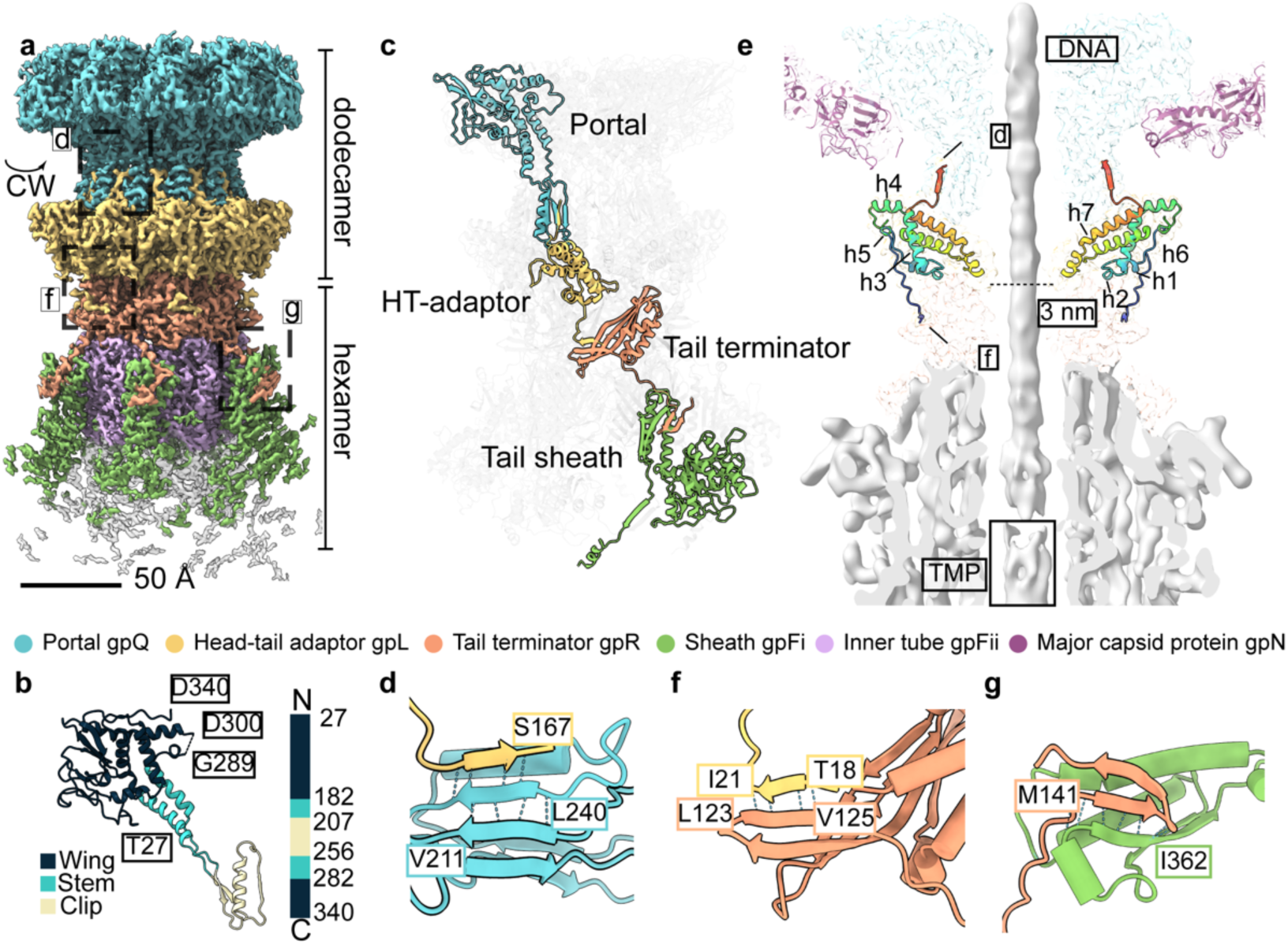
Reconstruction and model of P2 neck. (a) C6 local reconstruction of the P2 neck (PDB ID 29NE / EMD-57255; contour level 0.45), highlighting key interactions within the complex. (b) Monomer of the portal protein coloured based on domain organization. (c) Representative interaction modules illustrating the portal– HT-adaptor interface, the HT-adaptor-tail terminator interface, and the tail terminator-sheath interface. (d) Handshake antiparallel β-strand interactions between the portal and connector (e) Vertical cross-section of the symmetry-mismatched portal-capsid vertex reconstruction revealing lumenal density. The head-tail adaptor is coloured by sequence, highlighting its core helical bundle and the two terminator handshake motifs. (f–g), Handshake antiparallel β-strand interactions between (f) the connector and tail terminator, and (g) the tail terminator and sheath.

The tail of P2 continues from the neck towards the baseplate. The tail adopts the classical helical assembly of myoviruses, with the contractile sheath assembled around the rigid inner tube (Fig. 1a). We analysed the pre-contraction state, where the 192 copies of the sheath protein gpFi wrap around the inner tube in a right-handed C6 helical symmetry (rise = 39.5 Å, angle = 18°, diameter = 19.4 nm, Fig. 1a). The gpFi subunits of one ring interact with those of adjacent rings via interlocking terminal domains that form characteristic “handshake” interfaces found in well-described myophages and tailocins^13,37,42,43^. The inner tube, formed by gpFii exhibit a helical geometry (rise = 39.5 Å, angle = 18°, diameter = 4 nm). Post-contraction, the sheath is disengaged from the inner tube and shortened to approximately 41% of its extended length, preserving a helical C6 symmetry (rise = 16.4 Å, angle = 33°, diameter = 236 Å, Fig. 1d-e). Post-contraction, the N-terminal loop of gpFi undergoes a rotation of approximately 91°, while the C-terminal domain rotates by ∼21° (Fig. 1e).

The tail ends towards the baseplate of P2, which is minimal compared to that of other myophages and contractile injection systems (Fig. 1a, Supplementary Fig. 4). At the core of the baseplate, there are three copies of gpD, forming the hub complex. The hub connects the baseplate to the inner tube via six copies of the tube initiator protein gpU, which sits atop a trimeric assembly of the hub.

The peripheral region of the baseplate is assembled from six copies of the glue protein gpX and six wedge complexes. Each wedge is a heterotrimer composed of two copies of gpJ and one copy of gpI. The C-terminus of gpI connects to the trimeric tail fibre protein gpH, forming the distal attachment point. Beneath the central hub (gpD), there is the trimeric spike composed of gpV.

### A unique head-to-tail adaptor mediates portal-tail symmetry transition

The cryo-EM reconstruction revealed the architecture of the neck-portal and its binding to the sheath and tube. We reconstructed two cryo-EM maps of this region, one masking the portal, and other masking the sheath-terminator region, both imposing C6 symmetry, and achieving a global resolution of ∼2.6 Å (Figure 2, Supplementary Fig. 3).

The portal complex is composed of twelve copies of gpQ. Foldseek searches of the refined atomic model for gpQ find the portal proteins of phage Rcc01684 (PDB ID: 6TBA) and phage HK97 (PDB ID: 8FQL) as the closest portal proteins (Supplementary Fig. 4). P2’s portal protein is organized into three main regions: wing, steam and clip (Fig 2b). Differently from the portal proteins of Rcc01684 and HK97, P2 does not have a crown domain (Supplementary Fig. 4).

The clip domain facilitates the connection to the head-to-adaptor (HT-adaptor) protein gpL, located beneath the portal, via antiparallel β strand interaction between gpQ residues L240– Y243 and the C-terminal region of gpL (‘handshake’ interaction, Fig 2 c,d). Simultaneously, these residues form a parallel β-strand with adjacent gpQ subunit. Using Foldseek searches against the PDB, we found no structurally similar viral proteins, suggesting that gpL represents a novel fold, resembling phage adaptor proteins^15,44^. However, unlike other known adaptor proteins, gpL is characterized by a seven-helix core bundle connected by short, disordered linkers (Fig. 2e). The HT-adaptor presents a charged dipole inside its lumen, with negatively charged residues upwards to the head, and positively charged residues downwards to the tail. A similar charge profile is also seen in T4 adaptor protein, however, only its upmost region is negatively charged^44^.

The N-terminus of the HT-adaptor interacts with the clip domain of the portal, while its C-terminus (residues T18–I21, Fig. 2f) connects selectively to the tail terminator protein gpR in only six of the twelve copies. Density corresponding to the N-terminal residues of the HT-adaptor was missing: the first 17 residues were unresolved in HT-adaptor copies connecting to tail terminator gpR, and the first 27 residues were missing in the remaining six copies not directly contacting gpR. This asymmetric interaction is responsible for the transition from C12 to C6 symmetry at the neck.

The tail terminator gpR bridges the neck to the contractile sheath. gpR connects to the HT-adaptor via residues L123–V125 and anchors the sheath (gpFi) through an antiparallel β strand formed between gpR residues E142–I145 and the N-terminal domain of the apical sheath protein (Fig. 2g).

To resolve the internal architecture, symmetry was reduced by masking around a single C5 capsid vertex, enabling reconstruction of a C5-symmetrized capsid vertex map at 3.5 Å resolution (Fig. 2e). Subsequent relaxation of C5 symmetry by marginalization yielded a C1 reconstruction at 5.6 Å resolution. Low-pass filtering of this map to 8 Å revealed a discontinuity in the interior density at the neck-tail junction, consistent with the presence of packaged DNA and the TMP (Fig. 2e).

### The P2 tail terminator shares homology with siphoviruses

The uniqueness of P2 connector and tail terminator proteins prompted us to bioinformatically explore the relationships of these proteins and their homologs. Tail terminator shares similarities with the terminator proteins of other phages, as well as the cap proteins of extracellular contractile injection systems (eCIS) and tailocins (Supplementary Fig. 5).

Comparison of the tail terminator with the collar protein of R2 pyocin (PA0615), identifies the presence of the adaptor-terminator interaction, The tail terminator interacts with the sheath protein in a cascade of ‘handshakes’, which is fundamentally different than the pyocin cap-sheath interaction, that is anchored to different sheath subunit (Supplementary Fig. 5a).

Clustering of selected sequences and phylogenetic tree analysis show that the closest homologs of the phage P2 terminator protein are the terminators found in siphoviruses. Other myophages and contractile systems have distant sequence homology. Interestingly, the regions responsible for the handshake connections with the connector protein and the sheath are only conserved among phages in the *Peduoviridae* family (Supplementary Fig. 5b-c).

### P2 baseplate hub-tail initiator subcomplex breaks the C6 symmetry

Two localized cryo-EM maps of the baseplate, focusing on the hub and wedge were obtained at global resolutions of 2.83 Å and 3 Å, respectively (Fig. 3a, Supplementary Fig. 3). These reconstructions allowed us to confidently build atomic models for eight proteins within the baseplate region and to determine that many of the protein–protein interfaces are mediated through flexible or disordered loop regions.

**Figure 3.**
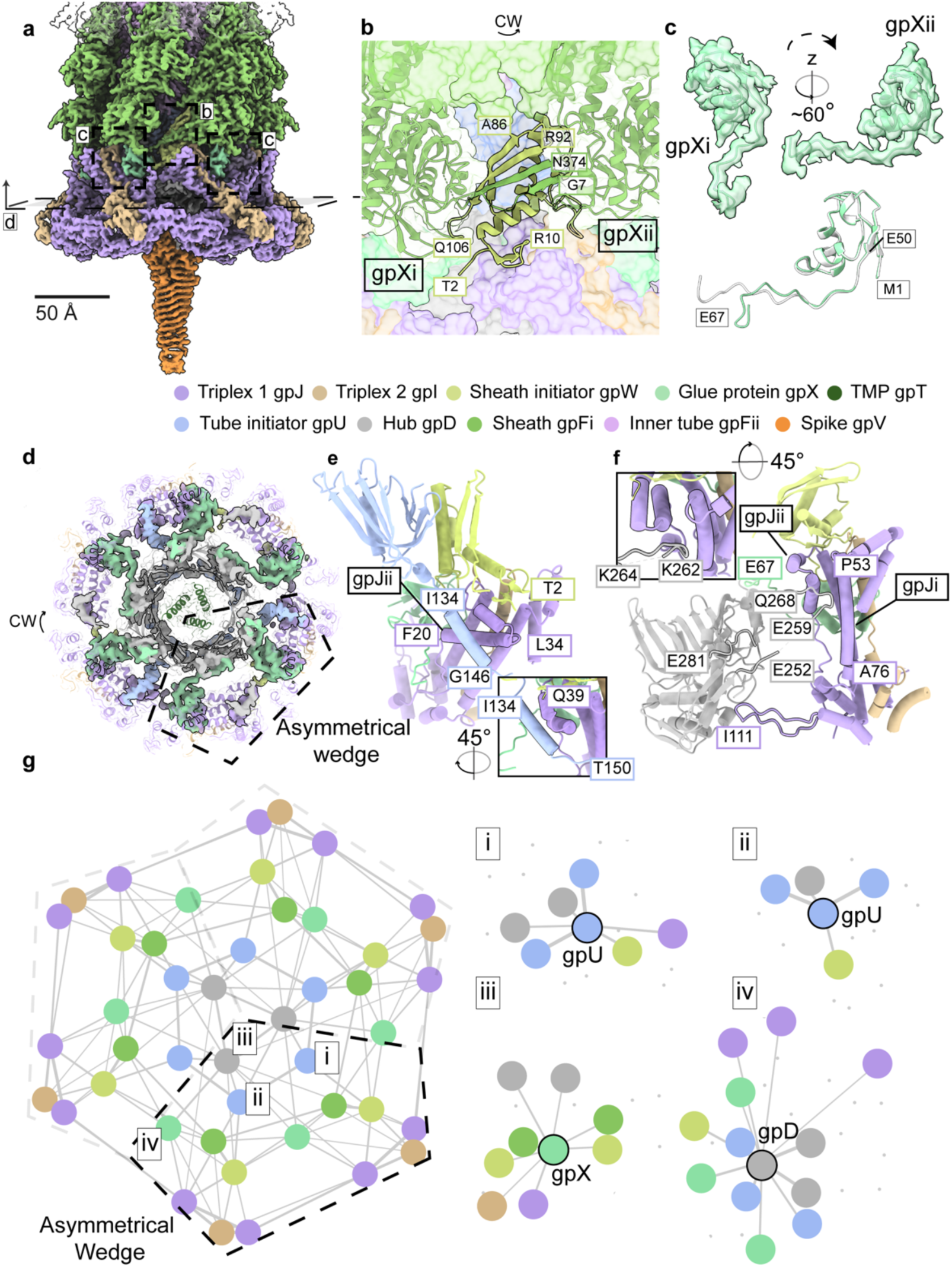
Reconstruction and modelling of P2 baseplate show it is a C3 complex, which is stabilized by minimal hub-tail initiator interactions. (a) C3 local reconstruction of the P2 baseplate wedge (PBD ID 29ND / EMD-57254; contour level 0.1), with regions corresponding to panels b–d indicated. (b) Model of the sheath initiator–sheath interaction (PDB ID 29ND). (c) Model of the glue protein, highlighting its extended disordered terminus. (d) Horizontal cross-section of the P2 baseplate hub map (contour level 0.1), showing the asymmetric unit. (e–f) Models of the extended tail tube initiator–triplex (e) and hub–triplex (f) close interface residues; (g) Interchain contact network generated using ProtCNet, identifying Cα contacts within an 8 Å radius. Asymmetric contacts formed by gpU and the symmetric contacts mediated by gpD are highlighted. CW: clockwise direction.

The baseplate is connected to the tail sheath via the sheath initiator protein gpW, which mimics the β-sheet “handshake” observed between adjacent tail sheath protein subunits along the sheath trunk (Fig. 3b).

The main interaction of the baseplate with the sheath occurs between the sheath initiator and its clockwise gpFi neighbour, involving the C-terminal region of gpFi (residues Asn374 to Thr382). Together, these interactions form a four-stranded β-sheet structure (Fig. 3b). This sheet includes the aforementioned β-strand of the clockwise gpFi, a parallel β-strand formed by the N-terminus of the counterclockwise gpFi subunit (residues Val10 to Gly7), and two β-strands contributed by gpW (residues Val78 to Ala86 and Arg92 to His101).

Located at the core of the baseplate wedge, gpW also contacts most baseplate proteins. gpW has a helix-loop-helix domain with three α-helices between residues Glu20 and Trp74. The loop formed by residues Asp52 until Pro57, contacts the clockwise gpFi, the loop located within Pro33 and Leu47 contacts the clockwise glue protein (gpX) and the triplex components gpJ and gpI, whereas the N-terminal disordered loop contacts the hetero trimer of gpJ/gpI, and the counterclockwise gpX.

Two alternative conformations of the glue protein (gpXi and gpXii) were modelled within the asymmetric unit, both spanning residues Met1 to Glu67. Interestingly, gpXii is tilted along the z-axis by approximately 60° when compared to gpXi (Fig. 3b). The N-terminal region of the glue protein adopts a canonical LysM fold, a domain typically located at the C-terminus of several eCIS tail initiator proteins (Supplementary Fig. 4) ^31,43,45^. The C-terminal portion of the P2 glue protein (from residue Glu50 onward) adopts two distinct conformations. In one wedge, the glue protein folds toward the triplex complex, whereas in the adjacent wedge it bends toward the hub (Fig. 3b-c).

The inner tube is connected to the baseplate through the tube initiator protein gpU. By symmetry relaxation, we resolved two distinct conformations of gpU. Together, these conformations assemble into the tube-initiator hexamer that links the hub to the inner tube in a C3 manner. We modelled the hub and spike proteins under the same symmetry.

The hub-tube initiator complex is responsible for breaking the C6 symmetry of the baseplate (Fig. 3d). Within one side of the asymmetric unit of the baseplate, we confidently modelled the tube initiator up to residue T150, whereas on the bordering side the density becomes untraceable beyond residue A129. In the longer model, we identified a C-terminal α-helix that engages the triplex complex through an electrostatic interaction (Fig. 3e). On the adjacent triplex complex, the hub mediates a similar electrostatic contact (Fig. 3f), together forming an asymmetric baseplate contact wedge (Fig. 3g). As this architectural feature has not been observed in previously described myophages or other contractile systems, we further investigated the uniqueness of the hub-initiator complex.

### P2 gpD and gpU asymmetric interaction with the baseplate wedge is a key feature of *Peduoviruses*

The P2 hub exhibits C3 symmetry and comprises three copies of hub protein gpD and six copies of tail initiator protein gpU. GpU is positioned toward the tail tube, whereas gpD interfaces the spike protein. GpD shares its core fold with hub proteins from other structurally characterized phages and eCIS (Supplementary Fig. 6). In P2, the gpD hub–triplex contact loop sterically interferes with the three gpU initiator–triplex contact helices, producing a highly distinctive architectural arrangement compared with other phages and eCIS (Fig. 3, Supplementary Fig. 6-7). We could only model the C-terminal helix of the tail initiator in three of the six copies. To assess whether this gpD–gpU symmetry break configuration is conserved across related virions, we examined the distribution of the gpD hub-loop and the gpU initiator-helix across phages. We queried gpD, gpU, and the major capsid protein (used as a proxy for *Peduoviruses*) against the NCBI-viruses database using BLAST. Sequence hits were clustered using MMseqs, and structures of cluster representatives were predicted with ESMFold. We manually annotated elongated loops at the gpD-wedge interface and extended helices in gpU. Two loop-extension variants (L12 and L23) were identified in gpD predictions (Supplementary Fig. 6a).

Among the 27 phages encoding a P2-like major capsid protein together with identifiable gpD and gpU homologs, 19 encoded both the loop and the helix, 5 encoded only the helix, and 3 encoded neither feature. In contrast, among 18 phages lacking a P2-like major capsid protein but encoding gpD- and gpU-like homologs, none encoded both features simultaneously: only two possessed the helix alone, one possessed the loop alone, and the remaining fifteen encoded neither.

To test if the loop and helix co-occur more frequently than expected under independence within P2-like phages, we performed Fisher’s exact test on the presence/absence matrix. The two features showed a significant positive association (one-sided p ≈ 0.02; φ ≈ 0.54), consistent with a functionally coupled structural module that is largely conserved. A second Fisher’s exact test comparing P2 major capsid-containing phages versus non-P2 major capsid-containing phages revealed strong enrichment of the full module in P2-like systems (19/27 vs 0/18; one-sided p ≈ 9 × 10⁻⁷). These findings indicate that the loop-helix pair represents a P2-specific architectural innovation rather than a broadly conserved feature among phages encoding gpD- and gpU-like proteins.

### P2 tail fibre is held by gpI flexible C-terminal region, which breaks local symmetry

Tail fibres adopt dynamic conformations, with a predominant stable state retracted toward the sheath. Local refinement of a subset of baseplate particles using a wedge-focused mask enabled reconstruction of a low-resolution model of the upward fibre conformation (Fig. 4a; Supplementary Fig. 3). Two distinct classes of the fibre-adaptor region were identified, differing by a ∼6° angular offset (Supplementary Fig. 8). Reconstruction of the full-length fibre for each class revealed distinct anchoring interfaces with the sheath: class 1 contacts the sixth sheath layer at the distal anchor domain (RBPseg-D82, Fig. 4b), whereas class 2 engages the third and fourth sheath layers (Supplementary Fig. 8). Class 1 fibres yielded higher-quality density and were used for subsequent analyses.

**Figure 4.**
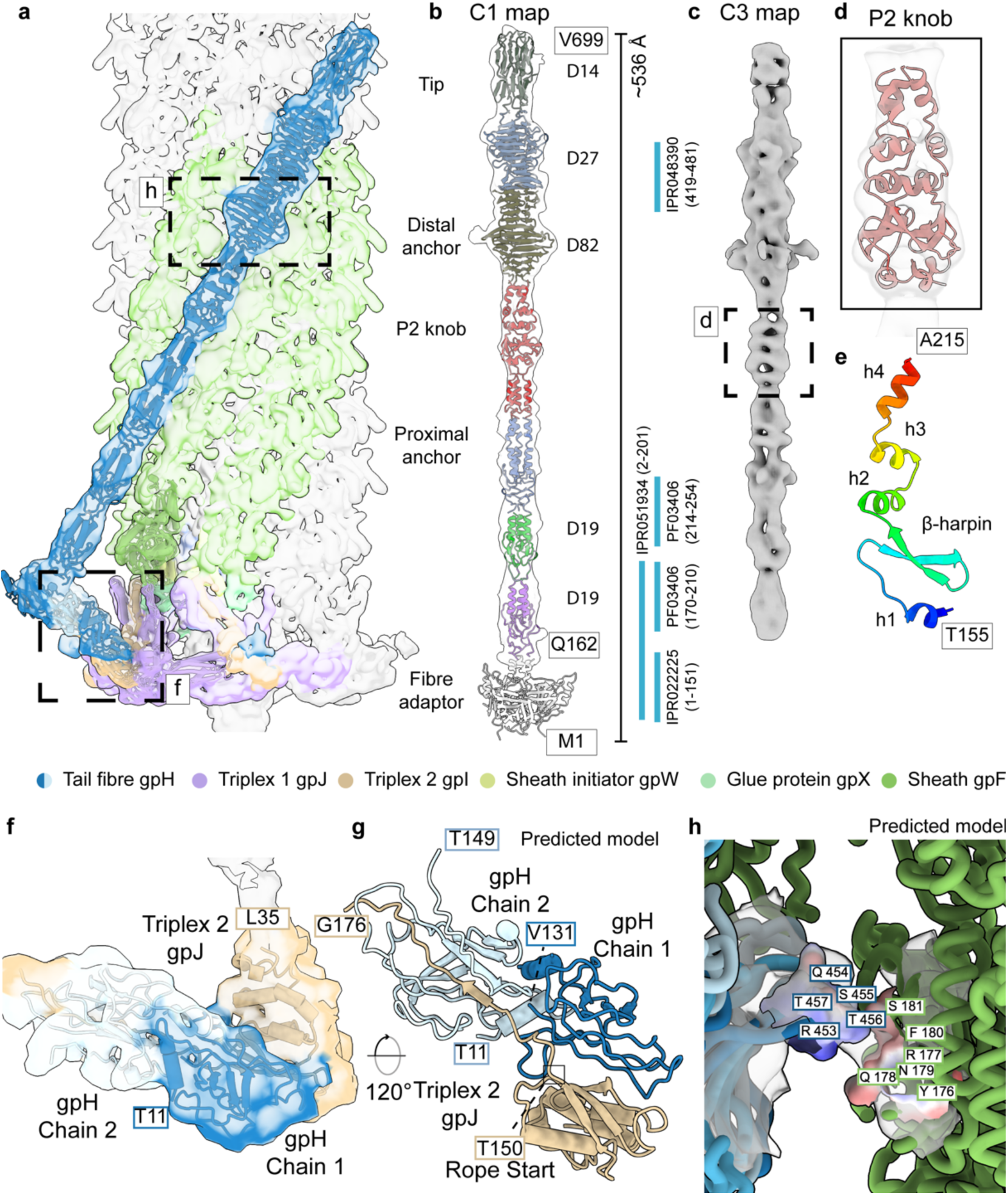
Low-resolution reconstruction of the P2 fibre supports computational predictions of a rope-like adaptor mechanism and anchoring domain. (a) Locally refined map of the P2 fibre (EMDB: EMD-57256) with superimposed RBPseg models (AF3-derived segments) merged from residue Q162 upward, together with refined adaptor predictions for the gpI–gpH dimer and gpH monomer. Third fibre-adaptor chain is only added as reference. (b) C1 map from (a) with RBPseg model of the P2 tail fibre, with pseudo-domains defined using RBPseg-sDp. Domain boundaries were annotated using InterPro^51^ and RBPseg-classify. (c) C3 local refined map of P2 fibre. (d) Superimposition of refined prediction for P2 knob domain with map in (c). (e) Single chain of P2 knob domain coloured based on amino acid sequence showing key structural features. (f) Local side view of the manually refined gpI rope-domain prediction, including the anchor adaptor. (g) Bottom view of the predicted fibre adaptor-gpI rope architecture. (h) Contact-density map of the fibre anchor domain, highlighting predicted interacting residues and corresponding electrostatic interfaces.

The fibre is a homotrimer of gpH, whose structure was predicted using the RBPseg v1.1.3 pipeline in combination with Alphafold3^46^ and fitted into the cryo-EM density (Fig. 4a–b). The P2 fibre is approximately 54 nm long and comprises eight domains: an N-terminal adaptor domain, two repeat domains (RBPseg-D19), two extension domains including the proximal anchor region, a distal anchor domain (RBPseg-D82), a β-helical knob domain – also found in proximal tail fibres of *Straboviridae* phages (IPR048390/RBPseg-D27) – and a distal fibre tip (RBPseg-D14; Fig. 4b). The distal tip adopts the same overall fold as those described for phage Mu and the short tail fibres of phages Bas49 and Bas54^20,47,48^. A C3 local refined map of P2’s fibre was used to confidently fit and refine a predicted model for the domain in between the distal and proximal anchor, which could not be bioinformatically annotated (Fig. 4c-d). We call it the P2 knob domain. The P2 knob domain is characterized by four short helices and one β-harpin motif (Fig. 4e).

The adaptor domain engages the baseplate through β-sheet interactions between two gpH subunits and the triplex protein gpI, forming a rope-like interface (Fig. 4f-g). Initial fitting of the predicted adaptor structure was suboptimal, suggesting local disruption of C3 symmetry upon gpI binding. Although gpI was partially built in the baseplate model, its C-terminal region (from residue R153) lacked interpretable density. To further characterize this interface, gpI-gpH interactions were modelled using Alphafold2-multimer with varying gpH stoichiometries (Supplementary Fig. 8). In models containing two or three gpH subunits, gpI consistently localized between two gpH chains with low predicted alignment error, resembling the rope fibre architectures described on diverse phages^42,49,50^. While these predictions could not be uniquely fitted into the low-resolution density, two chains from a rope fibre structure (PDB 8JOU) aligned well with both adaptor maps and were used as a template to guide placement of the Alphafold2 predictions (Fig. 4f-g, Supplementary Fig. 8). Differently from previously described fibre ropes, the P2’s rope is a domain of gpH, and not an additional protein.

Models from both adaptor classes fitted equally well into both wedges of baseplate asymmetric unit, indicating that the ∼6° angular offset does not arise from asymmetry within the baseplate core, but only from the different anchoring possibilities. One of the classes is anchored to the sheath via electrostatic interactions mediated by sheath residues Tyr176–Ser181. Although resolution was insufficient to assign specific contacting residues on the fibre, RBPseg predictions indicate likely contacts within the loop spanning residues Arg453–Thr457 (Fig. 4h).

### P2 tail measure protein follows the C3 spike symmetry

We reconstructed a C3-symmetrized cryo-EM map of the P2 spike gpV at a global resolution of 2.9 Å (Supplementary Fig. 9a). Although the local resolution decreases toward the apex domain, we could unambiguously visualize the characteristic iron-coordination site^52,53^ (Supplementary Fig. 9b). Toward the C terminus of the spike protein, we resolved three α-helices, into which we could confidently build a model for the C-terminal region of the TMP (Supplementary Fig. 9c).

To obtain more detailed structural information on the P2 TMP, we repositioned the box centre upward along the spike map and re-extracted particles using a larger box size. This reconstruction revealed a low-resolution density corresponding to the TMP (Supplementary Fig. 9d,e). The locally refined maps revealed elongated, helical-like bundle densities. However, the resolution was insufficient to allow confident atomic modelling.

Closer inspection of the internal density suggests that the TMP follows the C3 symmetry from the spike. Still, certain map slices appear consistent with the presence of six chains, as has been proposed for other myophages^13^. The oligomeric state of TMPs remains an active area of investigation, with recent studies supporting a six-chain organization often in a C3 arrangement^13,18,42^. Our data also suggest that P2’s TMP does not assemble into a canonical C6 coiled-coil bundle. Instead, the TMP may form two sets of spatially separated C3 chains, either stacked along the longitudinal axis or interlaced with one another.

### Key structural components on *Peduovirus* and *Peduoviridae*

We used BLASTP searches against all *Uroviricota* proteins in NCBI, first to identify homologs of bacteriophage P2 structural and orthogroup proteins. Proteins were grouped by their NCBI reference identifiers, and *Peduoviridae*-like phages were selected based on the presence of the complete set of P2 orthogroups. This dataset enabled a systematic assessment of the conservation of P2-like phage structural components relative to phage P2 (Supplementary Fig. 10a). Secondly, hierarchical clustering based on pairwise sequence identity of the major capsid protein was performed to visualize phylogenetic relationships among these viruses.

Notably, all members of the genus *Peduovirus* encode the full complement of 15 structural proteins with relatively high sequence similarity, although the tail fibre protein gpH is the least conserved. This overall conservation indicates that the P2 virion architecture is highly representative of the *Peduovirus* genus. The tail initiator-hub loop L23 and the C-terminal helices are sequence-conserved among *Peduovirus* members and are also present in phages belonging to the genera *Vimunumvirus* and *Duonihilunusvirus* (Supplementary Fig. 10b).

Genes encoding the capsid and neck proteins retain conserved genomic loci across all phages examined. The same conservation of gene order is observed for tail and baseplate components; however, in comparison with the R2 pyocin, the gene encoding the glue protein is located in different genomic regions. The tail fibre protein and TMP represent the most diverse structural proteins within this group. Notably, *Gegevirus* ST437OXA245phi41 lacks genes encoding tail and baseplate proteins, with the exception of the glue protein.

The genomic position of the glue protein gene is conserved among all P2 phages analysed but differs in the R2 pyocin, where it is located between the hub and the tail initiator (ripcord) genes. This altered gene context in the tailocin may indicate why, in eCIS systems, the tail initiator protein carries a LysM domain at its C-terminus.

In *Peduoviridae* phages, the TMP is encoded upstream of the tube initiator and hub genes, a genomic organization also observed in R2 pyocin, where the tube initiator is replaced by the ripcord protein. Given the predicted helical nature of both TMP and ripcord, we tested whether ripcord shares distant sequence homology with the C-terminal region of the P2 TMP. Homologous sequences were identified using HHblits, followed by sensitive profile–profile comparisons using HHpred/HHalign. These analyses did not reveal statistically supported sequence similarity, indicating that any shared evolutionary origin is not detectable at the sequence level.

### P2 tail pierces the outer membrane, and post-contracted particles lack baseplate integrity

We reconstructed cryo-EM maps of the neck and tail of post-contraction P2 particles (Fig. 1). However, we did not detect any discernible density corresponding to the post-contraction baseplate in any of our datasets (Supplementary Fig. 2, Fig. 5), despite extensive 2D and 3D classification. This absence may reflect substantial flexibility of the wedge region, or complete disassembly of the baseplate upon sheath contraction. To investigate this further we established two strategies for generating contracted P2 particles: (1) naturally triggered contraction with adsorption to WT host, (2) naturally triggered contraction with P2 adsorption to minicells, followed by membrane disruption with chloroform. The first approach was analysed by cryo-electron tomography (cryo-ET), and negative-staining TEM (NS-TEM). The sample concentration was insufficient for high-quality cryo-EM data collection in any of the approaches.

**Figure 5.**
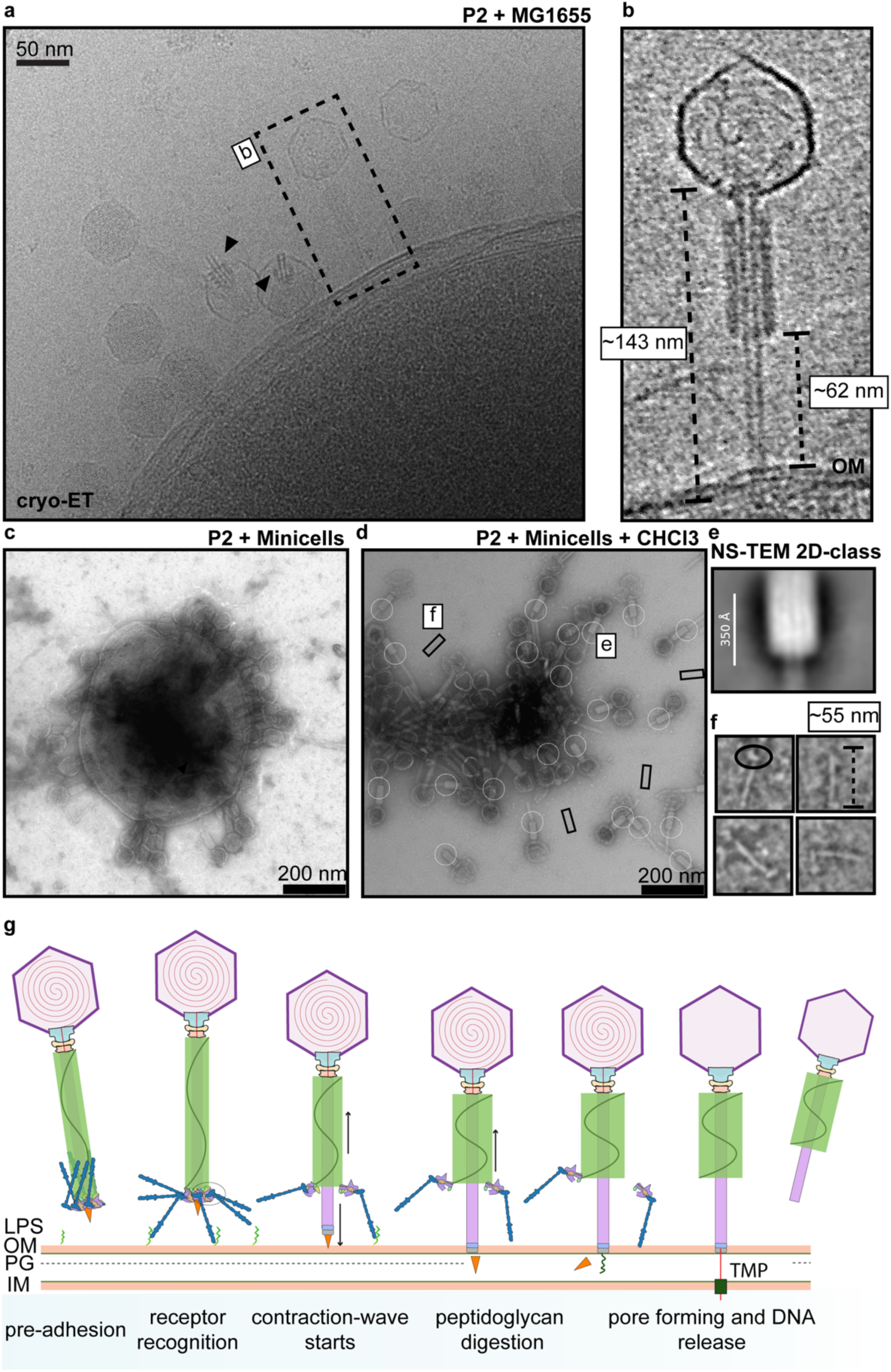
*In situ* analysis of P2 adsorption and proposed mechanism of DNA translocation. (a) Tilt series micrograph of P2*vir* interacting with *E. coli* MG1655. Black arrows indicate non-adsorbed post-contraction particles. (b) Cropped visualization of reconstructed tomogram of post-contraction P2*vir* particle adsorbed to *E. coli* MG1655. (c-f) Negative staining TEM (NS-TEM) micrograph of P2*vir* in different conditions: (c) adsorbed to minicells and (d) minicell samples after CHCl_3_ addition. (e) 2D class average of highlighted (circle) particles on (d). (f) zoom in of squared highlighted particles on (d), black circle highlights the probable baseplate triplex complex (g) proposed mechanism of DNA delivery. Phage P2 uses its lateral tail fibre complex to interact with the lipopolysaccharide core, which triggers the contraction wave. P2 spike is used to digest the peptidoglycan, allowing the release of the TMP. We propose the TMP forms a pore-like structure that allows DNA to cross the inner membrane. At the post-contraction state, the distance of the sheath and the outer membrane is bigger than the tail fibre length. Strongly bound fibres can be ripped from the particle at the late contraction state. Tomography data suggests some particles might be released after contraction.

Using cryo-electron tomography, we identified a small number of tomograms containing P2 particles, most in the post-contraction state. In the tomogram shown, one P2 particle is visibly attached to the host, whereas two are unattached (Fig. 5a–b). The reconstruction reveals major structural components of the post-contraction particles, including the empty capsid, neck, contracted sheath and inner tube (Fig. 5b). Neither the baseplate nor the fibres were visible in the attached or unattached contracted particles. The position and length of the inner tube suggest that it penetrates the outer membrane but does not reach the inner membrane (Fig. 5b). This observation suggests that attachment after contraction is primarily mediated by the inner tube–tube initiator complex. In the attached particle, the distance between the membrane-proximal end of the contracted sheath and the outer membrane is approximately 62 nm, exceeding the length of the P2 fibre. This geometry suggests that the fibre cannot remain attached to both the sheath and the outer membrane in the observed post-contraction state without substantial extension or rearrangement of its attachment complex. The increased separation may therefore promote detachment of the fibre–triplex complex as contraction proceeds.

The minicell-derived samples revealed similar results, with no clear baseplate around the sheath, and the presence of flexible and broken inner tubes (Fig. 5c). Following chloroform treatment, minicells were lysed and post-contraction P2 particles were released (Fig. 5d). Manual particle picking at the sheath-tube interface followed by 2D classification did not reveal any baseplate-like features (Fig. 5e). This agrees with the cryo-EM 2D-classes and 3D-reconstruction obtained in our original dataset (Fig. 1). Moreover, we observed fibre-like filaments with lengths of approximately 55 nm, which we hypothesize correspond to the fibre-triplex complex (Fig. 5f).

Together, these observations support the hypothesis that the P2 baseplate is no longer attached to the virion in the post-contracted infective state. The absence of the baseplate from the contracted phage is likely due to the loss of stabilizing interactions between the baseplate and the hub-tail initiator complex, as supported by our cryo-EM results, as well as the polling force between the contracting sheath and the host outer membrane.

Our cryo-ET observations suggest that the P2 inner tube does not reach the inner membrane, implying that an additional conduit is required for DNA translocation. To explore whether the TMP could fulfil this role after release from the tail tube, we generated trimeric and hexameric AlphaFold models. The hexameric model adopted a pore-like arrangement with a predicted channel diameter of approximately 3 nm, whereas the trimeric model formed an incomplete pore-like assembly (Supplementary Fig. 11). Transmembrane helix predictions identified five candidate helices between residues L211 and I612; in the hexameric model, the proposed membrane-spanning region extends approximately 5 nm (Supplementary Fig. 11). These predictions support the hypothesis that the released TMP could form a conduit across the inner membrane, a mechanism similar to the mechanism recently proposed for bacteriophage T4^54^.

Our structural and *in situ* observations suggest a mechanism linking receptor recognition, sheath contraction and DNA delivery (Fig. 5g). We propose that binding of the extended fibres to the lipopolysaccharide core, as previously described^23,55^, transmits a pulling force through the fibre–triplex complex, disrupting its contacts with the hub–tube initiator subcomplex and the glue protein to initiate contraction. Receptor-bound fibres may guide tube insertion while maintaining a connection between the sheath and the host surface. As contraction proceeds, disruption of stabilizing baseplate contacts, together with increasing tension across the fibres, may promote detachment of the fibre–triplex complex. This interpretation is consistent with the approximately 62 nm sheath–outer membrane separation observed in the attached contracted particle, which exceeds the approximately 55 nm length of the putative detached fibre–triplex complexes. Loss of this tether could reduce the mechanical coupling between sheath contraction and tube insertion, potentially limiting penetration depth. Following outer membrane penetration, the P2 spike may enter the periplasm, consistent with recent biochemical evidence placing gpV in the periplasm after contraction^56^. We further propose that the P2 spike may possess peptidoglycan-degrading activity, analogous to that of the T4 spike-associated gp5 protein^57^, although this activity remains to be experimentally demonstrated in P2. This allows TMP release from the inner tube. Because the tube appears not to reach the inner membrane, we propose that the released TMP assembles into an additional membrane-spanning conduit for DNA translocation, consistent with its predicted pore-like architecture and candidate transmembrane helices. The sequence of these events and the proposed roles of fibre detachment and TMP pore formation remain to be experimentally established.

## Conclusions

We present the near-complete structure of *Peduovirus* P2 in both pre- and post-contraction states. Phage P2 possesses a T=7 dextro icosahedral capsid and a distinctive C6 neck assembled from the portal, connector, and tail terminator proteins. The P2 connector protein belongs to a conserved orthogroup within the *Peduoviridae* family; here, we demonstrate that it is not only sequence-divergent but also lacks structural similarity to previously characterized phage neck components. In contrast, the tail terminator protein is homologous to the cap protein found in tailocins and shares features with terminator proteins from siphophages.

The P2 tail is composed of the tape measure protein (TMP), sheath, and inner tube proteins and closely resembles previously described contractile systems, such as the R2 pyocin. Cryo-EM density corresponding to the TMP is observed within the lumen of the inner tube. Toward the distal spike, the TMP adopts a three-helix bundle conformation. Further along the tail, the resolution is insufficient for model building, and the oligomeric state of the TMP remains unresolved. Nevertheless, cross-sections through the TMP density reveal clear C3 features, with possible six chains. This is supported by AlphaFold predictions of an hexameric pore-like structure.

We show that P2 harbours a minimal baseplate compared to other contractile injection systems. The tube initiator and tail hub constitute defining features of the P2 baseplate and together break the symmetry of the structure from C6 to C3. Their complex stabilizes the wedge in concert with the sheath initiator and the glue proteins. Using structural bioinformatics, we identify that the C-terminal helix of the tube initiator and the L23 loop of the hub are key features of P2 phages, suggesting that these elements have co-evolved, and define a P2-specific mechanism.

Low-resolution maps of the P2 tail fibres reveal symmetry breaking around the fibre adaptor region, which likely adopts a rope-like fold mediated by the gpI extension acting as an attachment tether. Although resolution was not sufficient for modelling on the extension rope, this hypothesis is strengthened by *in-silico* analyses. Conformation heterogeneity of fibre’s upward retracted state, indicates the fibre can be anchored upwards in two different regions of the sheath, likely hold by electrostatic interactions, indicating that ionic gradient differences might promote fibres to extend. In P2-phages the fibre is the least conserved structural protein by sequence, which indicates its evolutionary plasticity to adapt to new hosts.

Together, our pre- and post-contraction structures and in situ observations support a model for P2 DNA translocation in which receptor recognition by the extended fibres initiates sheath contraction through disruption of stabilizing baseplate contacts. As contraction proceeds, increasing tension across receptor-bound fibres may promote detachment of the fibre–triplex complex from the sheath, potentially limiting the depth of tube penetration. Our observations further suggest that the inner tube penetrates the outer membrane but does not reach the inner membrane. Following spike-mediated peptidoglycan cleavage and TMP release, we propose that the TMP forms an additional conduit across the inner membrane to enable DNA translocation. This model links the minimal baseplate architecture of P2 to a proposed mechanism of fibre-mediated activation, baseplate disassembly and sequential penetration of the bacterial envelope.

## Methods

### Phage purification

A primary stock of phage P2*vir* was propagated and phage was purified using standard phage purification protocols. In summary, P2*vir* was first propagated using a small batch lysate (50 mL), where the host (*E. coli* MG1655 ΔRM) was grown until OD600 of 0.2 in LB media supplemented with 20 mM MgSO4 and 5 mM CaCl2, and infected with an MOI < 1 for 4-5 hours until OD600 dropped below 0.2 at 90 RPM. Lysate was treated with 1% (v:v) chloroform for 5 minutes with slow shaking, followed by 10 minutes of no shaking. Supernatant was saved and used to infect a 1L of host with same conditions. After OD600 dropped below 0.2, lysate was treated again with 1% chloroform, followed by addition of 5 μg/mL of DNase 1 and RNase A for 1 hour with low vortexing. After we added of 10 mM EDTA. Next, we did PEG precipitation overnight (30 g/L NaCl and 75 g/L PEG 8000 overnight at 4 °C with rotating magnet at 200 rpm). The phage was pellet at 15,000 x *g* for 60 minutes and resuspended in 5 mL of SM buffer (100 mM NaCl, 8mM MgCl, 50 mM Tris-HCl pH 7.5). We added chloroform 1:1 (v:v) in the resuspended phage, and inverted the sample until homogenized, next we centrifuged it at 6,000 x *g* for 15 minutes and collected supernatant. Next, we load the supernatant on an OptiPrep™(Sigma-Aldrich) gradient (50%-10%) and was centrifuged at 124,000 x *g* for 18 h. Bands with phage were collected, and dialyzed in 10 kDA Thermo Scientific Slide-A-Lyzer dialysis cassettes against SM buffer. Finally, P2 particles were concentrated at 20,000 x *g* for 20 minutes and resuspended in 50 μL of SM buffer.

### Cryo-electron microscopy data collection

Purified P2*vir* particles were deposited onto UltrAuFoil R2/2 grids coated with a 2-nm continuous carbon film and rapidly vitrified in liquid ethane cooled by liquid nitrogen using a Vitrobot Mark IV. Cryo-EM data were acquired on a Thermo Scientific Krios G2 microscope equipped with a Falcon 4i direct electron detector and a Selectris X energy filter, using a total electron dose of 42 e⁻/Å² and a physical pixel size of 1.2 Å.

### Cryo-electron microscopy processing

Unless stated otherwise, all cryo-EM data was processed using CryoSPARC v4.5.1^58^ and visualized using UCSF ChimeraX v1.6-v1.10^59^.

### Pre-processing

Two independent cryo-EM datasets were collected and processed separately. The first dataset (8,675 movies) was used to reconstruct the pre-contracted state, whereas the second dataset (21,540 movies) was used to reconstruct the post-contracted state. Both datasets were pre-processed following the same workflow. Beam-induced motion correction was performed using Patch Motion Correction, and contrast transfer function (CTF) parameters were estimated using Patch CTF Estimation, both with default settings. All exposures were manually inspected and curated prior to downstream processing.

### Initial particle picking and 2D-classes

Initial particles were automatically picked using the Blob Picker and inspected using the Inspect Particle Picks job. Picked particles were extracted with a box size of 256 px and subjected to 2D classification using default parameters. Classes corresponding to P2 tail particles were selected and used as templates for template-based particle picking, followed by re-extraction with a larger box size of 560 px and subsequent 2D classification. This iterative process was repeated twice until well-defined classes corresponding to the neck and baseplate regions were obtained.

### Baseplate reconstruction

Approximately 51,000 particles were used for an initial *ab initio* reconstruction, followed by two rounds of homogeneous refinement and a subsequent non-uniform refinement. The resulting map was used for template-based particle picking, followed by 2D classification to further clean the dataset. Particles selected for the final non-uniform refinement were symmetry-expanded (C6) and subjected to focused 3D classification using a mask around the spike region, with two classes, a target resolution of 3.5 Å, hard classification enabled, and a class similarity threshold of 0.8. This classification yielded two distinct classes that preserved the C3 symmetry of the baseplate.

For each class, focused masks were generated around the spike, core, and wedge regions, and C1 local refinements were performed. The resulting particle sets and maps were re-extracted, and duplicate particles were removed. The cleaned particle sets were then used for C3 local refinement of the core region, followed by C3 symmetry expansion and subsequent C1 local refinement of the wedge region.

Local sharpening of the spike apex region was performed using DeepEMhancer^60^. Prior to sharpening, the box size of the locally refined map was reduced using the relion_image_handler tool implemented in RELION v4.0^61^. DeepEMhancer was then applied to the cropped map using the wideTarget model.

### TMP reconstruction

TMP reconstruction was initiated from C3-symmetrized baseplate particles aligned around the central spike. Particles were translated upward using Volume Alignment Tools and re-extracted with a box size of 1,056 pixels, applying Fourier cropping to 560 pixels. A new volume was generated using homogeneous reconstruction, followed by C3-symmetrized homogeneous refinement. The density within the tail lumen was then used for local refinement (C3), and a 3D classification into two classes was performed to remove junk particles arising from the re-extraction procedure. Selected particles were subsequently re-extracted without cropping and locally refined (C3) using a mask focused on the lumen density. The resulting map was low-pass filtered to 8 Å. To target the upward region of the TMP, particles were translated an additional 372 pixels upward and re-extracted again with a 1,056-pixel box size and Fourier cropping to 560 pixels. The same reconstruction and refinement workflow was applied to obtain a low-pass filtered map of the upper TMP region.

### Fibre reconstruction

The C6 symmetry-expanded baseplate map was used as the starting point for three successive rounds of focused 3D classification, initially applying a mask around the wedge region, followed by masks encompassing the fibre N-terminal region and the full-length fibre. Selected particles were re-extracted using a larger box size (900 px), and the final reconstruction was obtained using the Homogeneous Refinement job. Additional local refinements targeting specific fibre domains were performed but did not result in further improvement of map quality. The N-terminal domain of the fibre was locally sharpened using DeepEMhancer with the wideTarget model, following the same procedure applied to the spike apex region.

### Neck reconstruction

From 2D classes, approximately 38,000 particles were selected for *ab initio* reconstruction, followed by heterogeneous refinement and subsequent non-uniform refinement. These particles were then used to train a Topaz particle picker, which was applied to the full dataset to select approximately 138,000 particles. The newly picked particles were subjected to 2D classification, and classes corresponding to the neck region were selected (∼45,000 particles) and refined using homogeneous and non-uniform refinement.

The resulting particles and maps were further processed by heterogeneous refinement to separate particles containing capsids from those without capsid density, splitting virions from tail-only particles. Particles corresponding to intact capsids were symmetry-expanded with C6 symmetry and subjected to focused local refinement using a mask around the neck wedge region, followed by 3D classification. After removal of duplicate particles, additional local refinements were performed under C6 symmetry using masks targeting the portal region or the sheath/terminator region.

### Capsid reconstruction

Neck particles containing the capsid were re-extracted using a larger box size (1,280 px), Fourier-cropped to 640 px and recentred, followed by homogeneous refinement. The resulting capsid map was used for template-based particle picking. Picked particles were subjected to 2D classification and *ab initio* reconstruction, followed by heterogeneous refinement into three classes with icosahedral symmetry applied. Particles corresponding to the class displaying well-defined DNA density were selected for further homogeneous and non-uniform refinement. For high-resolution analysis of the capsomer region, selected particles were re-extracted with a box size of 720 px, symmetry-expanded, and subjected to focused local refinement using a mask around individual capsomers.

### Reconstruction of P2 in post-contraction state

The post-contraction reconstruction workflow is summarized in Supplementary Fig. 2. Initial 2D classification identified contracted sheath particles, from which ∼10,000 particles were selected for *ab initio* reconstruction using a box size of 512 pixels with Fourier cropping to 360 pixels. Three C1 *ab initio* classes were generated, separating particles corresponding to the neck, sheath-only, and downward sheath–inner tube regions. Particles from each region were then subjected to heterogeneous refinement into two classes to further separate well-resolved particles from junk.

For the lower tail region, template-based picking was used to retrain particle picking independently for each region. Approximately 20,000 particles were re-extracted using a large box size (1,440 pixels) with Fourier cropping to 360 pixels. After removal of duplicate particles, ∼10,000 particles remained and were further processed by 2D classification, *ab initio* reconstruction, and heterogeneous refinement. From this dataset, 2,500 particles were selected and re-extracted with a 540-pixel box size (Fourier cropped to 360 pixels) centred on the sheath–tube interface. These particles were used for homogeneous reconstruction and refinement with C6 symmetry, followed by symmetry expansion around the sheath wedge and subsequent 3D classification.

### Model building and refinement

For all proteins except the tail tape measure protein (TMP) and the fibre, AlphaFold2^62^-predicted models were used as initial templates. Predicted models were manually docked into the corresponding cryo-EM maps using UCSF ChimeraX^59,63^. Biopython^64^ (bio.PDB) was used to adjust files when needed. Automated real-space refinement was performed using RELION/StarMap^61,65^, followed by interactive manual refinement in ISOLDE 1.6.0^66^, Coot-1.1^67^.Models were further validated and real-space refined using Phenix version 1.21rc1-5084-000 ^68^.

An initial model for the terminal helical region of the TMP was generated using ModelAngelo^69^ and subsequently manually refined and rebuilt in ISOLDE.

### Prediction of fibre structure

Fibre modelling was performed using a combination of structure prediction and segmentation approaches. An initial monomeric fibre model was generated using ESMFold v1^70^. The fibre was segmented into structural fractions using the RBPseg v1.1.3^20^ pipeline, with clustering performed using the k-means method (rbpseg-sdp). Structural models for individual fractions were predicted using AlphaFold3^46^ assuming trimeric assemblies. The full-length fiber mode; was constructed using the rbpseg-merge workflow with an additional relaxation step.

The fibre-adaptor-rope region was modelled by fitting the previously reported rope fibre structure (PDB ID 8JOU) into the cryo-EM density using UCSF ChimeraX. Monomeric Alphafold2 predictions of two gpH subunits and gpI were aligned to PDB ID 8JOU using the Matchmaker tool with default parameters and individually fitted into the corresponding low-resolution density using Fit-in-Map. The resulting composite model was used as input for Phenix Dock in Map with default settings. The docked model was subsequently refined by Amber relaxation in two stages: first with 2,500 steps and an energy tolerance of 0.1, followed by 5,000 steps and an energy tolerance of 0.5.

### TEM analysis of P2 with minicells

We constructed an *E. coli* MG1655(Seq) (CGSC Strain #7740) derived strain producing “skinny minicells”^71^ using CRISPR-Cas engineering based on previous protocol^72^(*E. coli* MG1655(Seq) *minCDE mreB*-A125V-2MM (bNTL054)). A 1 L LB culture of this strain was inoculated with 2 mL overnight culture and grown at 37 °C with shaking (180 rpm) to OD₆₀₀ ≈ 1.0. Parental cells were removed by sequential centrifugation at 2,000 × g and 3,500 × g for 5 min each at 4 °C, discarding the pellets. The supernatant was centrifuged at 15,000 × g for 45 min at 4 °C to pellet minicells, which were resuspended in 1.5 mL PBS. Minicells were further purified on a discontinuous 5–20% OptiPrep gradient (3.5 mL each of 5%, 10%, 15%, and 20%) prepared in SW28 tubes and centrifuged at 2,000 × g for 30 min at 4 °C. Gradients were fractionated into 1.5 mL fractions (11 total) and analysed by light microscopy and negative staining. Fractions 3 and 4 (counted from the top), which showed maximal minicell enrichment with minimal parental cell contamination, were pooled and filtered through a 0.45 μm PVDF membrane. Filtered fractions were centrifuged at 20,000 × g for 1 h at 4 °C. The purified minicell pellet was resuspended in 100 μL PBS or M9 medium. For phage incubation assays, MgCl₂ was added to a final concentration of 20 mM.

Phages were incubated with minicells at room temperature for 1h. After that, samples were prepared for negative staining. First, 5 μL of sample was deposited into a carbon film on copper mesh grid for 45 s, followed by three rounds of wash with SM buffer, and addition of 5 μL 1% uranyl acetate for 45 s, and blotting. Micrographs were captured using a Morgagni 268 equipped with a side-mounted MegaView III camera.

### Cryo-ET of P2-infected *E. coli* cells

An exponentially growing culture of *E. coli* MG1655 ΔRM (OD₆₀₀ = 0.3) was infected with bacteriophage P2 and incubated for 10 min. A 1.5 mL aliquot of the infected culture was pelleted by centrifugation at 10,000 × g for 1 min at 4 °C. The pellet was resuspended in 1.5 mL of ice-cold phage buffer (50 mM Tris-HCl, pH 8.0, 150 mM NaCl, 2 mM CaCl₂, 2 mM MgCl₂) and centrifuged again under the same conditions. The final pellet was resuspended in 150 µL of ice-cold phage buffer and maintained on ice.

Samples were applied to holey carbon copper grids (R1.2/1.3, 200 mesh; Quantifoil), blotted, and vitrified by plunge-freezing in liquid ethane using an FEI Vitrobot Mark IV. Tomographic tilt series were recorded over an angular range of −60° to +60° with 3° increments using a dose-symmetric tilting scheme under low-dose conditions. Data were acquired on a Titan Krios G2 transmission electron microscope operated at 300 kV, equipped with a Falcon 4i direct electron detector and a Selectris X energy filter (slit width 10 eV) using Tomo5 software. Each tilt series was collected at a nominal defocus of −5 µm and acquired as a movie of 50 frames in 3 fractions. The total cumulative dose per tilt series was 50 e⁻/Å², and the calibrated pixel size was 1.9 Å.

Movie frames were corrected for motion using MotionCorr3^73^ and assembled into drift-corrected stacks using IMOD^74^. The drift-corrected stacks were aligned using patch-track alignment in IMOD within the eTomo package. Rigid body fitting of the P2 particle models to the tomogram was done using ChimeraX.

### Structural comparison of P2 proteins

Structural homologues of P2 protein monomers were identified using Foldseek^75^. Predicted P2 protein structures were used as queries against the PDB100 database (release 20240101), using the 3Di/AA search mode. For each query, the top three hits ranked by Foldseek score were selected for subsequent structural comparison.

### Contact network of baseplates

The contact networks were calculated using ProtCNet v1, developed in this work to visualize protein contact patterns. ProtCNet accepts PDB or mmCIF inputs, parses atomic coordinates per chain using Biopython^64^, and computes contacts using a KD-tree–based nearest-neighbour search (SciPy^76^). Contacts were defined using Cα atoms and an 8.0 Å distance cutoff (command: protcnet_cli.py --input_file file.pdb --cutoff 8.0 --cluster_chains --edge_style rectangle). Inter-chain contacts were quantified per chain pair and visualized as an interactive network graph (NetworkX^77^ rendered with Sigma/ipysigma^78^; edge weights proportional to the number of contacts)

### Bioinformatical analysis

Homologous proteins of P2 were identified using BLASTp^79^ searches against the NCBI RefSeq viral protein database (accession date: 17 December 2024). All viral proteins from this dataset were used to construct a local BLAST database, and individual P2 proteins were queried to retrieve homologous sequences.

For phylogenetic analysis, homologous sequences were aligned using MAFFT v7.525 (2024/Mar/13)^80^ with the globalpair algorithm and iterative refinement (maximum 100 iterations), using all available CPU cores. Poorly aligned regions were removed using TrimAl v1.5.rev0 build[2024-05-27] ^81^ with a gap threshold of 0.25. Phylogenetic trees were inferred using IQ-TREE version 3.0.1^82^, with 2,000 ultrafast bootstrap replicates, and 1,000 SH-like approximate likelihood ratio tests, using all allocated computational threads.

Phylogenetic trees were visualized using pyCirclize for circular representations or Bio.Phylo (Biopython) for standard layouts. Multiple sequence alignments were visualized using pyMSAviz. Structural models of gpU and gpD homologs were predicted using ESMFold v1^70^.

## Data and Code Availability

The cryo-EM maps were deposited at EMDB. Entries: EMD-57344, EMD-57343, EMD-57258, EMD-57256, EMD-57341, EMD-57255, EMD-57254. Structures PDB ID: 29NY, 29SG, 29NE, 29ND.

ProtCNet pipeline is deposited at: 10.5281/zenodo.15440765

## Acknowledgments

The Novo Nordisk Foundation Center for Protein Research is supported financially by the Novo Nordisk Foundation (NNF14CC0001). N.M.I.T. acknowledges support from an NNF Hallas-Møller Emerging Investigator grant (NNF17OC0031006), an NNF Hallas-Møller Ascending Investigator grant (NNF23OC0081528) and an NNF Project grant (NNF21OC0071948) and is also a member of the Integrative Structural Biology Cluster (ISBUC) at the University of Copenhagen. V.K-S acknowledges the Novo Nordisk Foundation Copenhagen PhD Programme for grant NNF0069780. We acknowledge the Core Facility of Integrated Microscopy at University of Copenhagen (CFIM) for help with data collection. We acknowledge the Big Data Management Platform at Novo Nordisk Foundation Center for Protein Research for the computational resources.

We are thankful to Prof. Dr. Alexander Harms for sharing the P2*vir* isolate. We are thankful to Dr. Claudia S. Kielkopf for discussions meetings and supervision during project conceptualization, as well as for prof-reading the final manuscript.

## Author contribution

Conceptualization: V.K-S., A.R-E., N.M.I.T.; Methodology: V.K-S., A.R-E., M.S., M. Š., N.M.I.T.; Software: V.K-S.; Investigation: V.K-S., A.R-E., M.S., M. Š. Writing-original draft: V.K-S. Writing-review & editing: V.K-S., A.R-E., M.S., M.Š. and N.M.I.T.; Visualization: V.K-S., M. Š.; Supervision: N.M.I.T.; Funding acquisition: V.K-S. and N.M.I.T.

