## Supplementary figures and tables for "Piercing mechanism in the minimal contractile tail of bacteriophage P2"

974 **Supplementary Files**

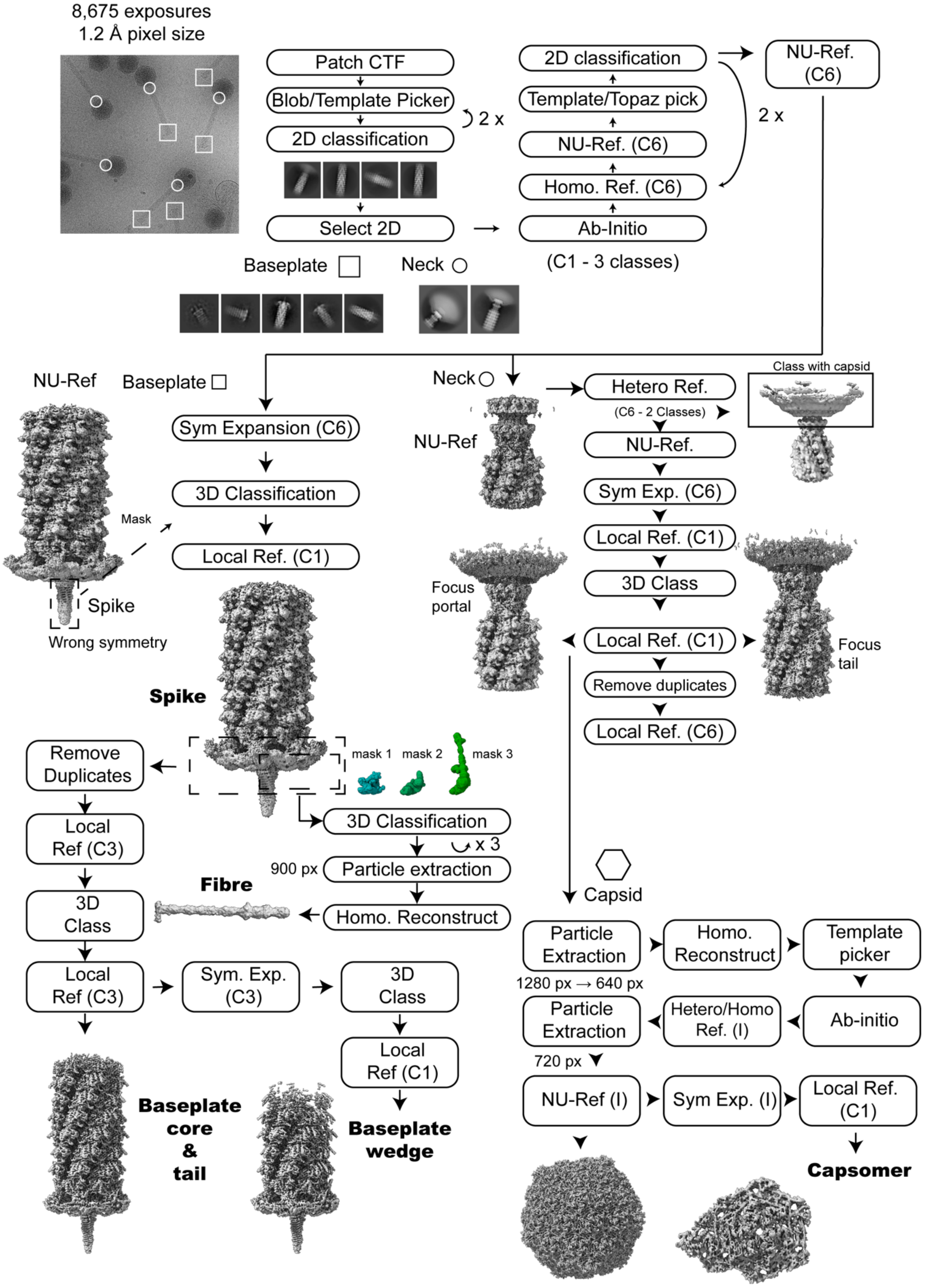


975

976 **Supplementary Figure 1. Cryo-EM processing pipeline for the pre-contracted P2 phage.**

A total of 8,765 micrographs were pre-processed followed by particle picking and 2D classification. Particle classes corresponding to the baseplate and neck were subjected to further refinement. The figure shows key intermediate reconstructions as well as the final deposited maps.

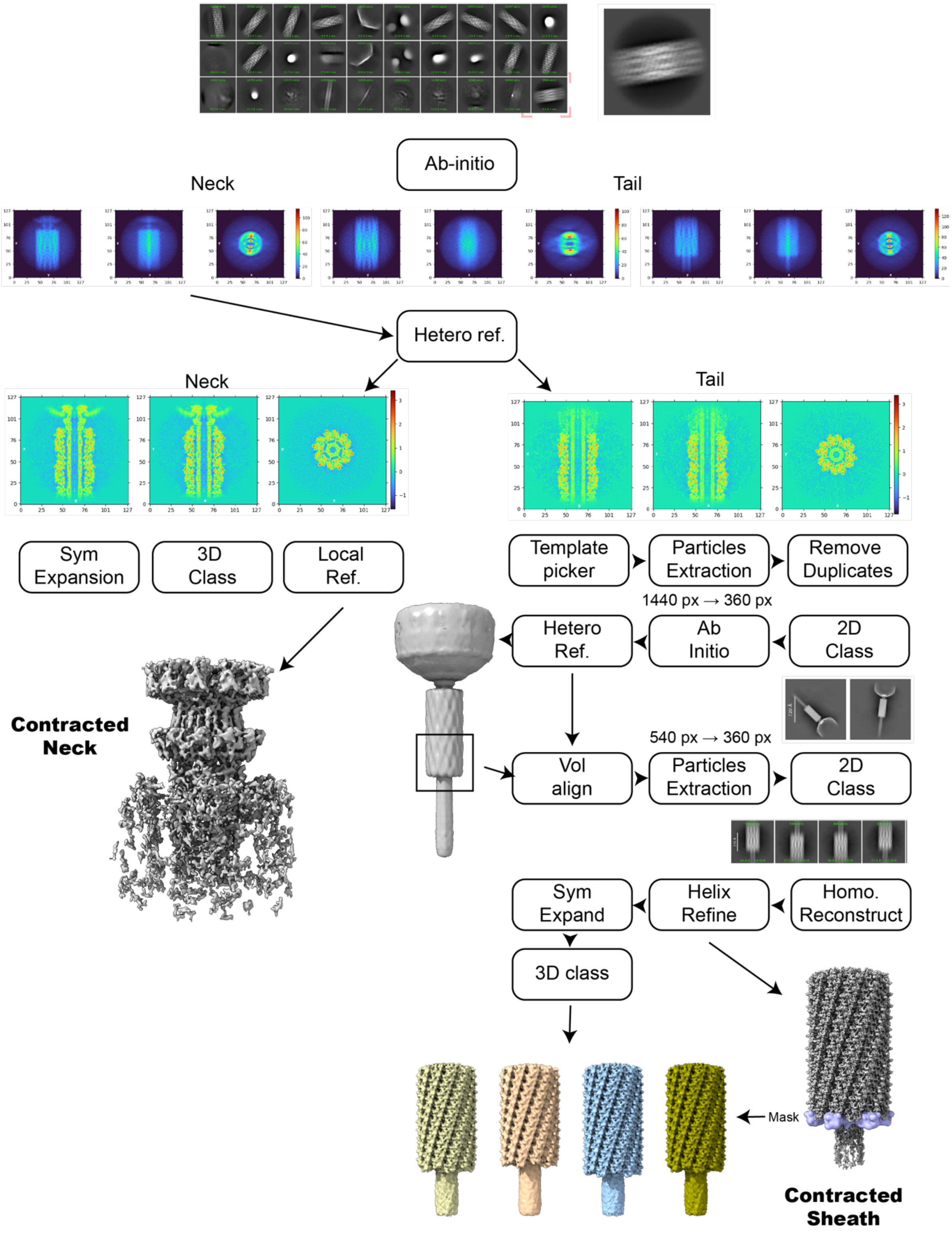

**Supplementary Figure 2. Cryo-EM processing pipeline of post-contracted phage.**

Micrographs were pre-processed, followed by particle picking and 2D classification. Classes corresponding to the post-contraction sheath were selected for further analysis. *Ab initio* reconstruction separated particles into two major regions: the apical–neck region and the base–sheath region. Extensive 3D classification was then performed to identify the baseplate-containing subset. The figure presents key intermediate reconstructions as well as the final deposited maps.

Baseplate Core

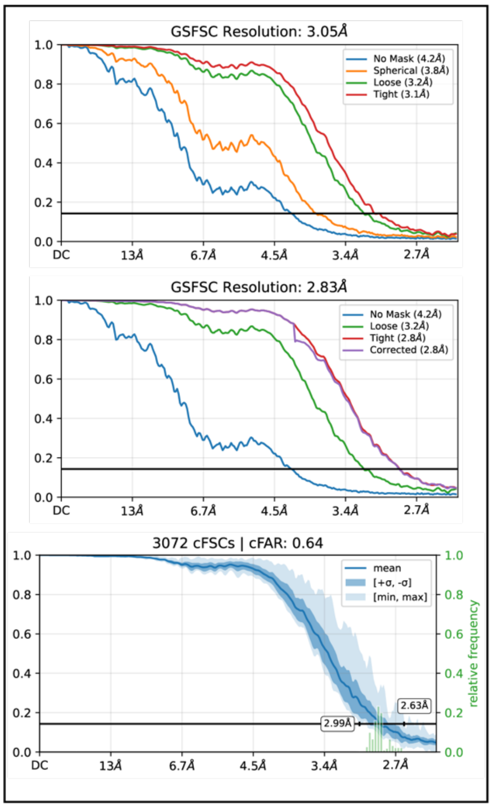


Wedge Core

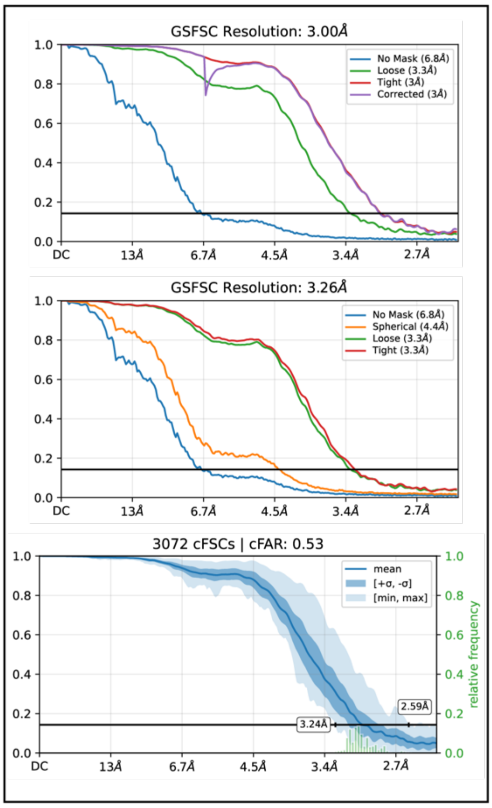


Neck Portal

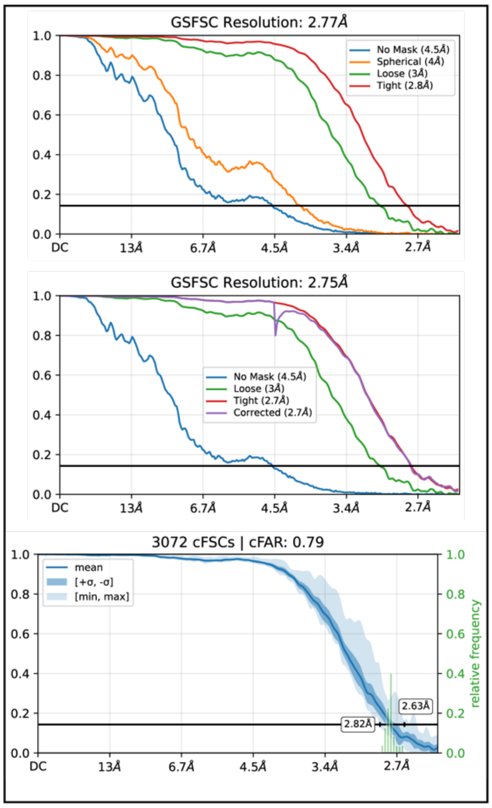


Neck tail

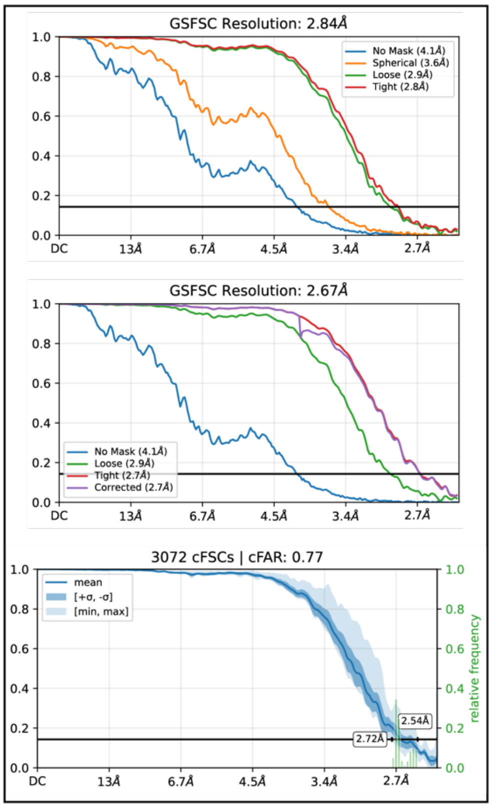

Capsomer

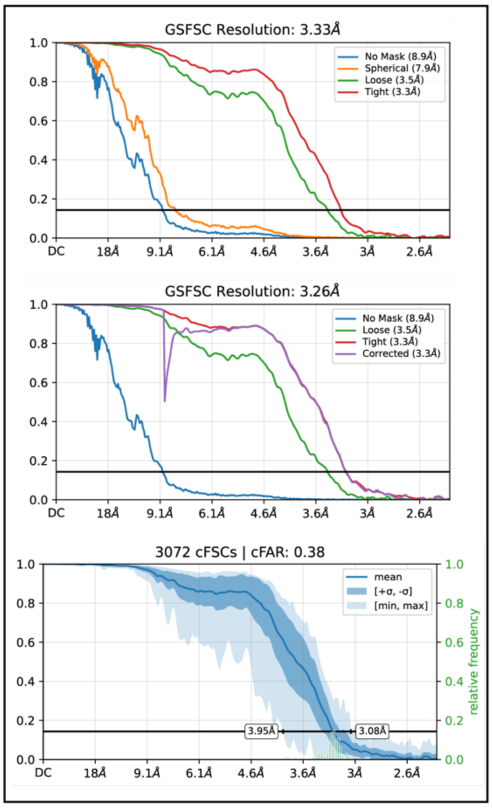


Tail Fibre

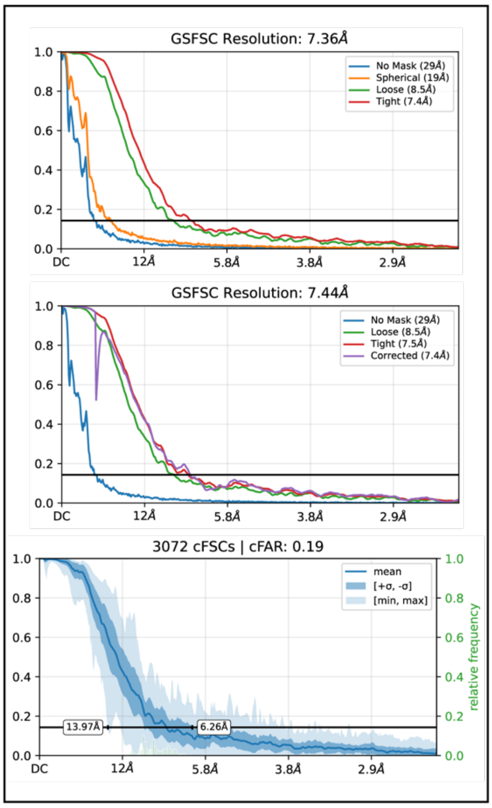


Baseplate-fibre rope adaptor

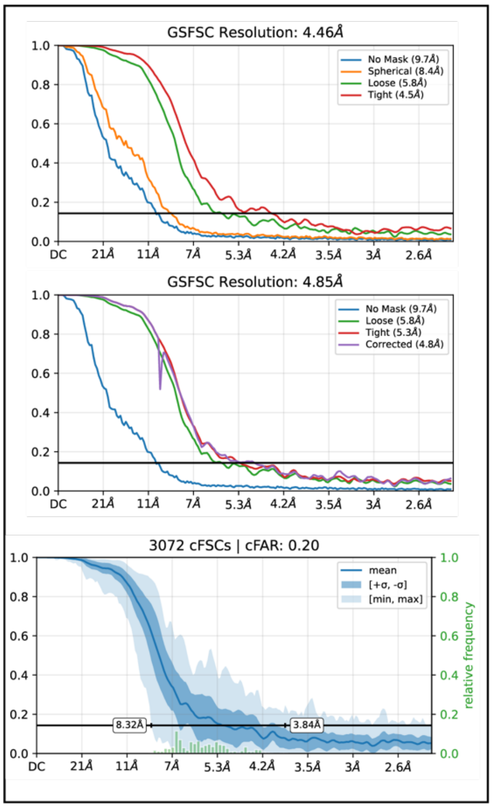


Post-contraction neck

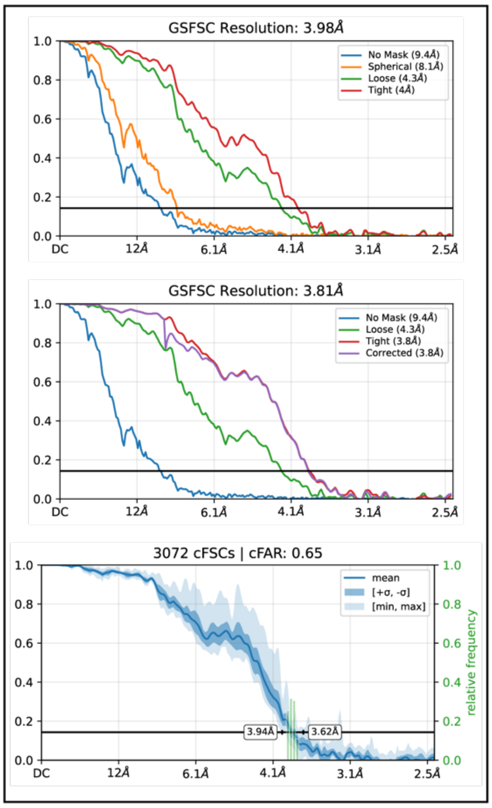

Post-contraction tail

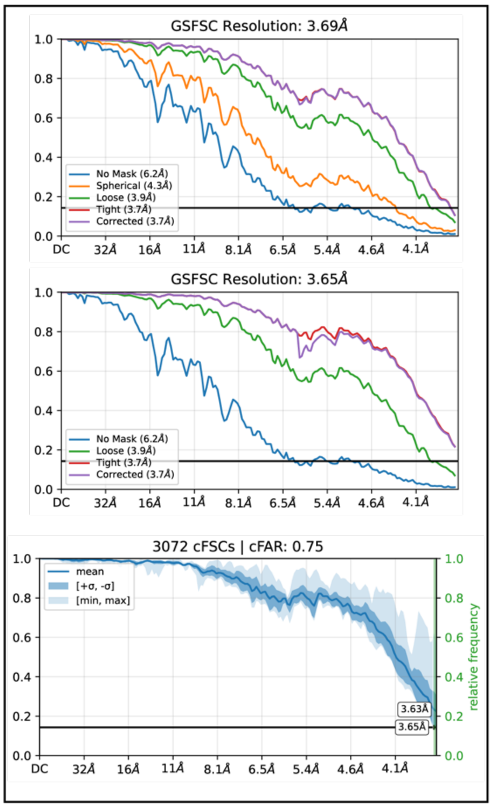


**Supplementary Figure 3. FSC curves and global resolution estimates**

Global Fourier shell correlation (GSFSC) and corrected FSC (cFSC) plots obtained from cryoSPARC refinement jobs for all maps in this manuscript. For each reconstruction, FSC curves are shown for unmasked and masked half-maps, including loose and tight masks, as well as corrected FSC curves where applicable. Reported global resolution values correspond to the FSC = 0.143 criterion following mask auto-tightening and correction.

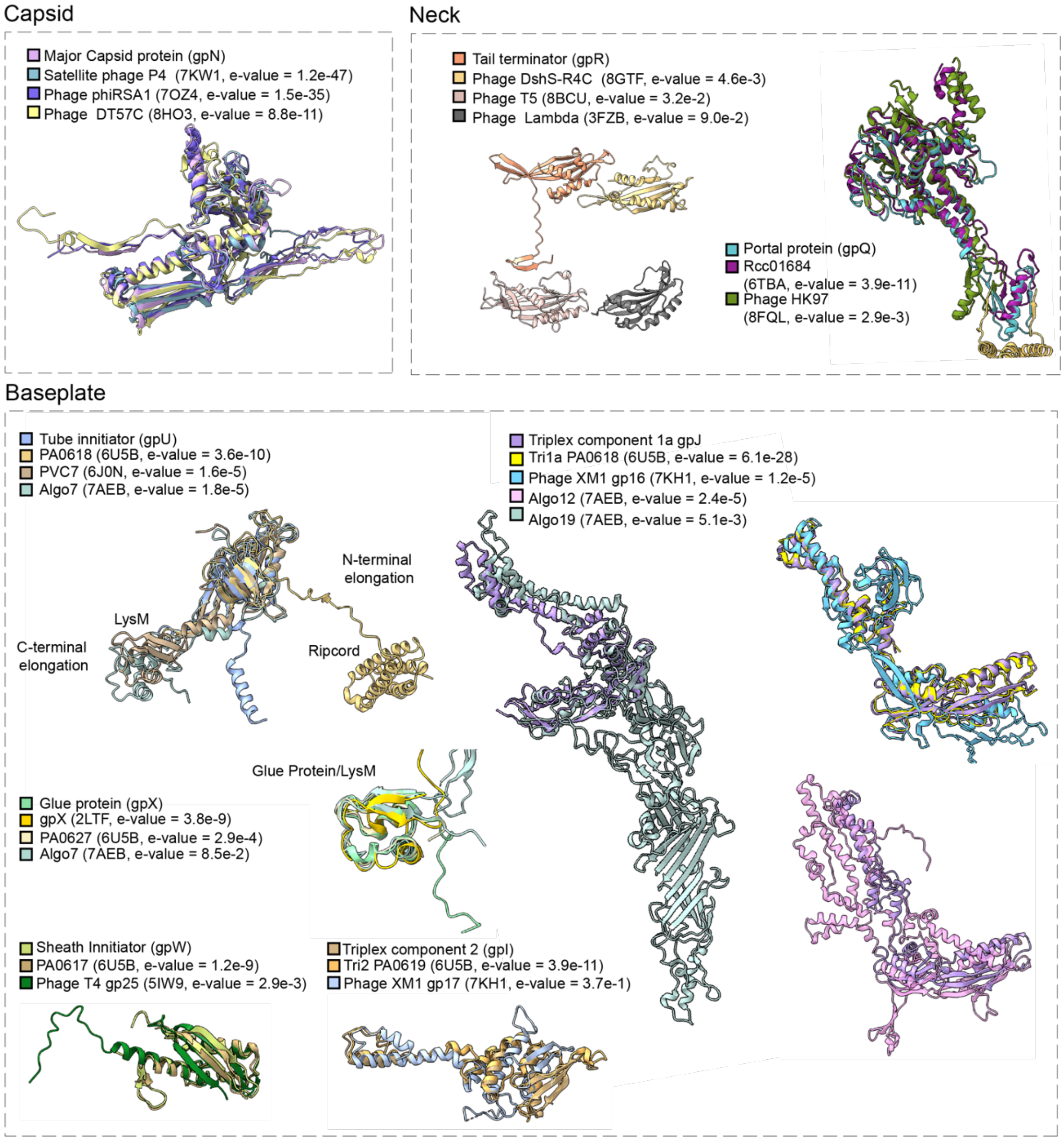


**Supplementary Figure 4. Structural comparison of P2 proteins with distant homologs**

Structural comparisons of P2 proteins from the capsid (top left), neck (top right), and baseplate (bottom) with their top structural matches identified using Foldseek against the PDB100 database (release 20240101). P2 protein models are shown superimposed with the corresponding Foldseek hits. Alignment significance is indicated by the Foldseek E-values, which are represented in the colour legend.

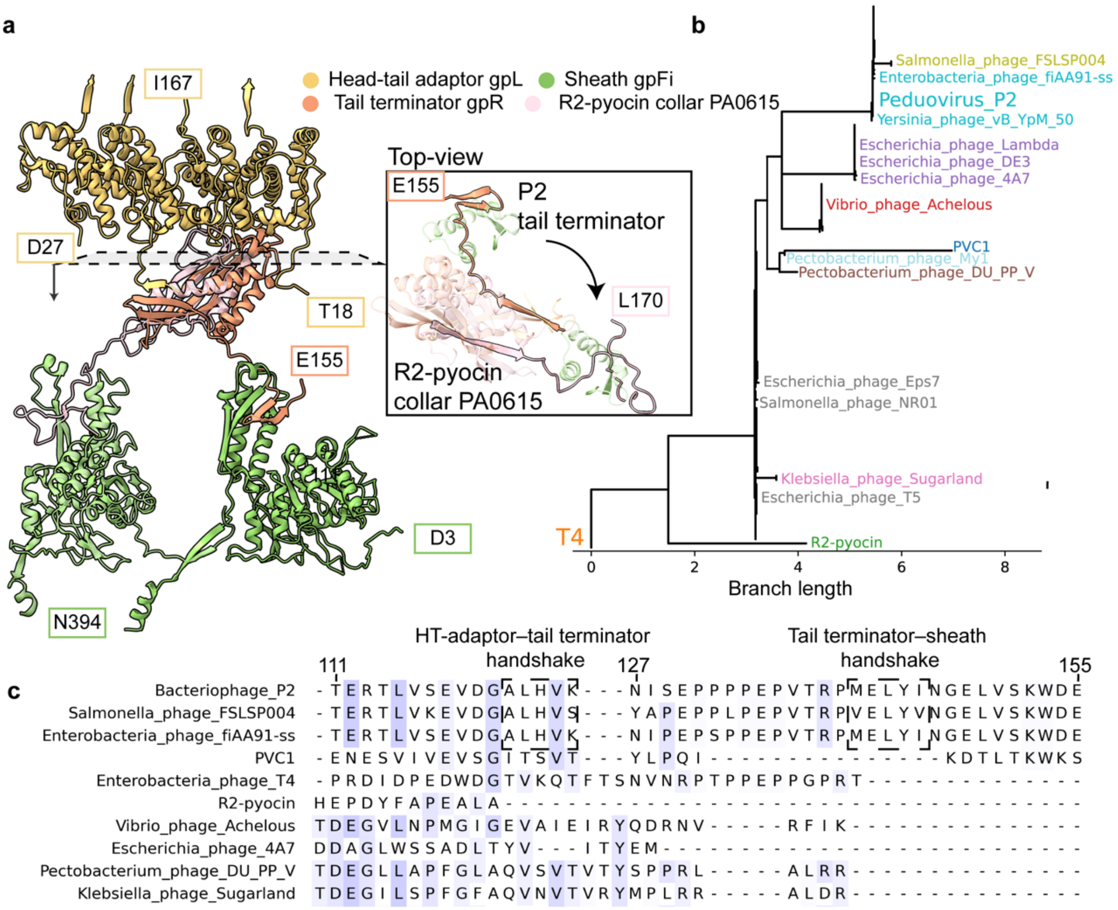


**Supplementary Figure 5. P2 neck shares distant homology with siphophages and contractile injection systems**

(a) Structural comparison of the P2 tail terminator–sheath complex (PDB ID: 29NE) with the R2-pyocin CAP–tail sheath complex (PDB ID 6U5F; coloured light pink and light green). (b) Phylogenetic tree constructed from the tail terminator sequences selected in panel (c), coloured according to clustering and rooted on the phage T4 tail terminator. (c) Multiple sequence alignment of selected tail terminator proteins from diverse phages and CAP proteins from PVC1 and R2 pyocin. Structural homologues were identified using Foldseek against the PDB100-20240101, and matched proteins were queried using BLAST against NCBI-viruses (download date: 20241217). Sequences were clustered with MMseqs , and representative sequences were selected for MSA. The alignment highlights the regions corresponding to handshake β-strand interactions.

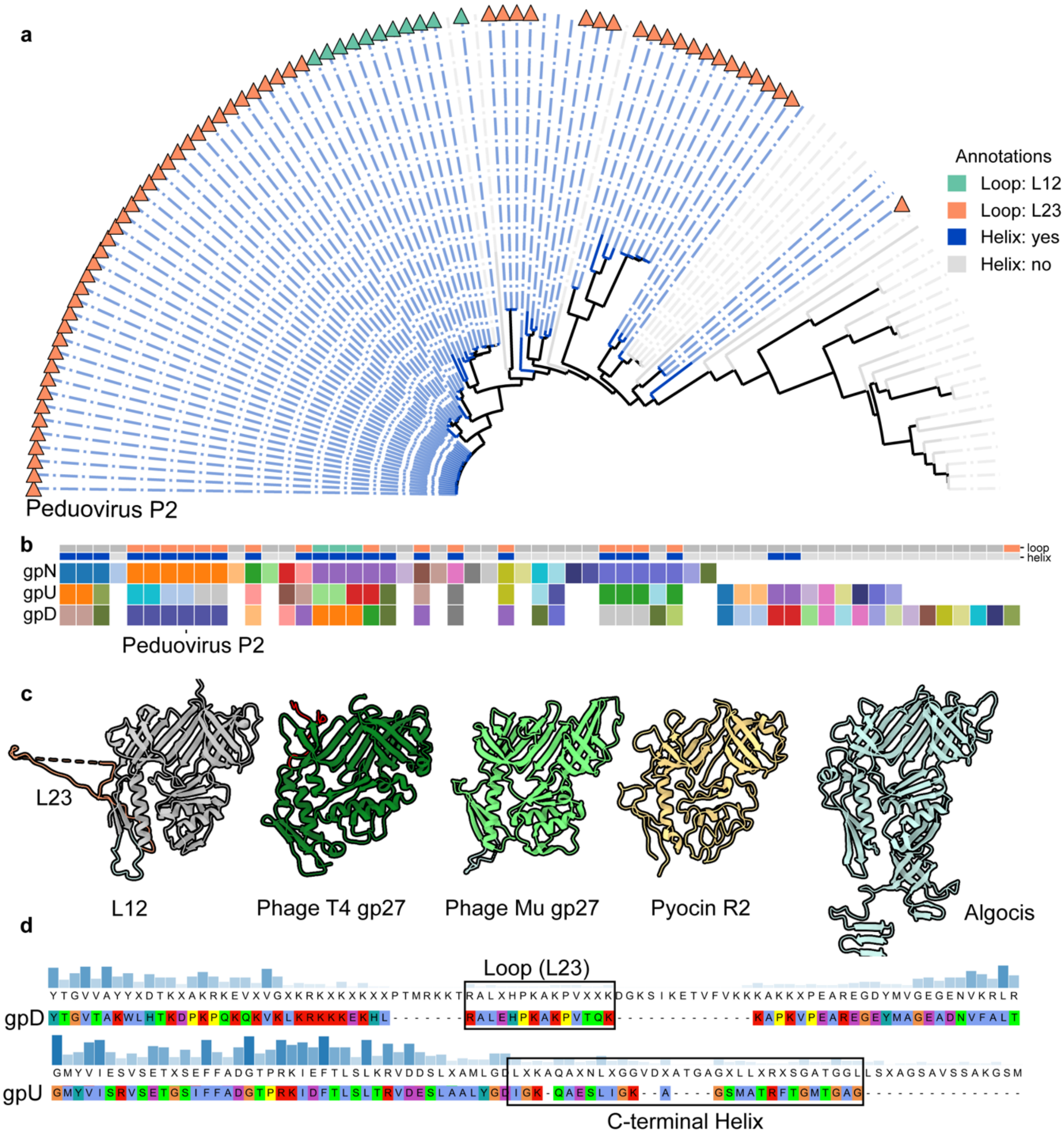


**Supplementary Figure 6. P2 hub has evolved two long loops that are commonly found on *Peduoviridae*.**

(a) Phylogenetic tree of NCBI viruses encoding homologs of both gpU and gpD, constructed using MAFFT based on gpN sequence similarity. Branches are colored according to the presence (blue) or absence (gray) of the gpU C-terminal helix in the cluster representative. The presence of extended loop insertions is indicated by triangles: loop L12 (green) and loop L23 (orange). (b) Overview of sequence representatives for NCBI viruses encoding homologs of gpN, gpU, or gpD. The top row indicates the presence or absence of extended gpD loops L12 or L23 (colored as in a). The second row denotes the presence (blue) or absence (gray) of the gpU C-terminal helix. The third, fourth, and fifth rows indicate structural clustering based on Foldseek analysis, colours represent the structural class. (c) Structural comparison of hub proteins from phage P2, T4 (PDB ID 5IV5), Mu (PDB ID 9KI1), pyocin R2 (PDB ID 6U5B), and AlgoCIS (PDB ID 7AEF). In the P2 hub, loops L12 and L23 are highlighted. (d) Sequence alignment of the P2 gpD loop and gpU C-terminal helix, colored according to the Clustal Omega scheme, illustrating sequence conservation relative to representative sequences shown in b.

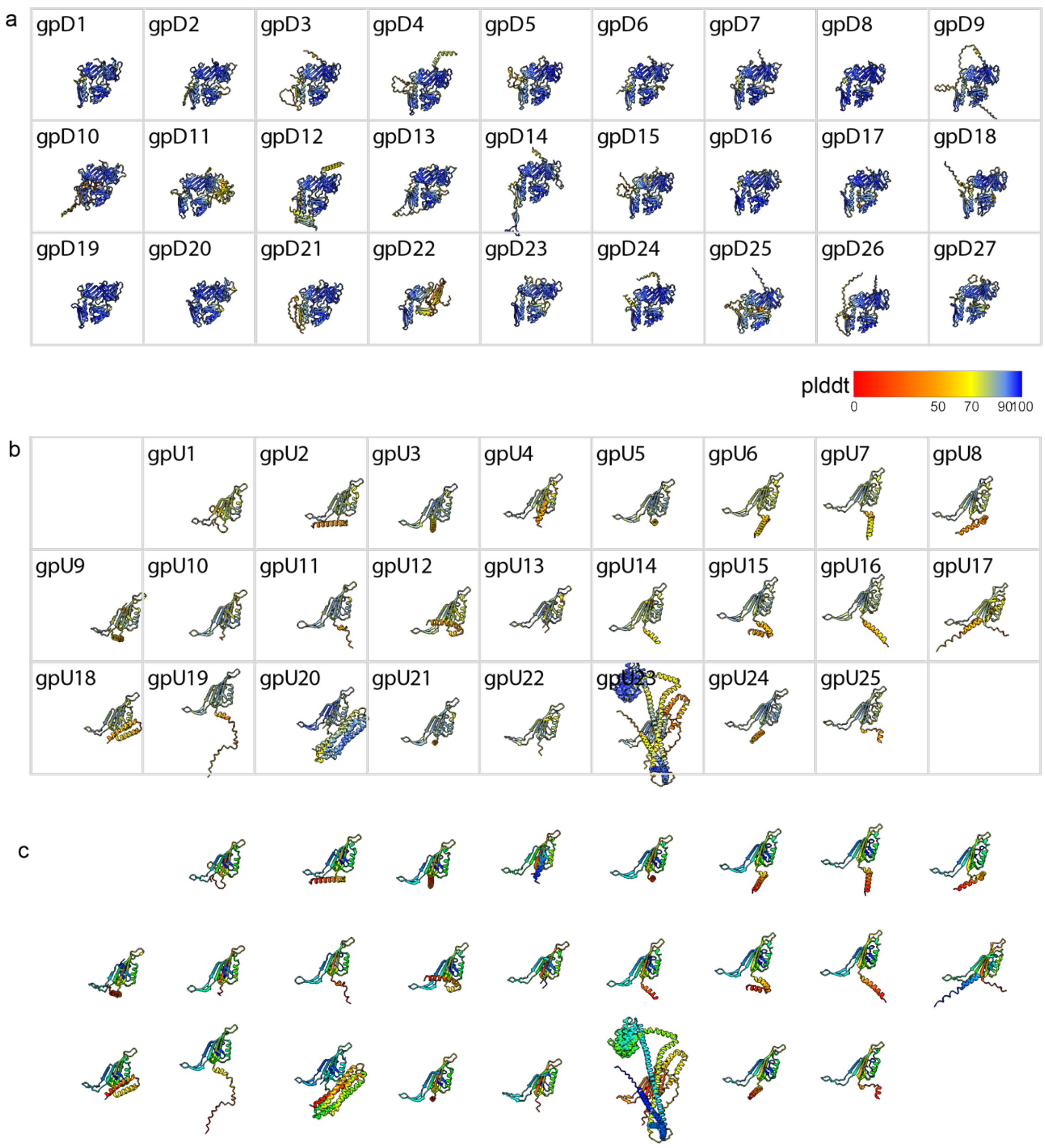


**Supplementary Figure 7. ESMfold models of gpU-D homologs and confidence metrics.** (a) ESMFold-predicted structures of gpD homologs. (b) ESMFold-predicted structures of gpU homologs. (c) Alternative visualization of gpU homologs. In a and b, structures are coloured by per-residue confidence (pLDDT). In c, structures are coloured by residue position using a rainbow gradient.

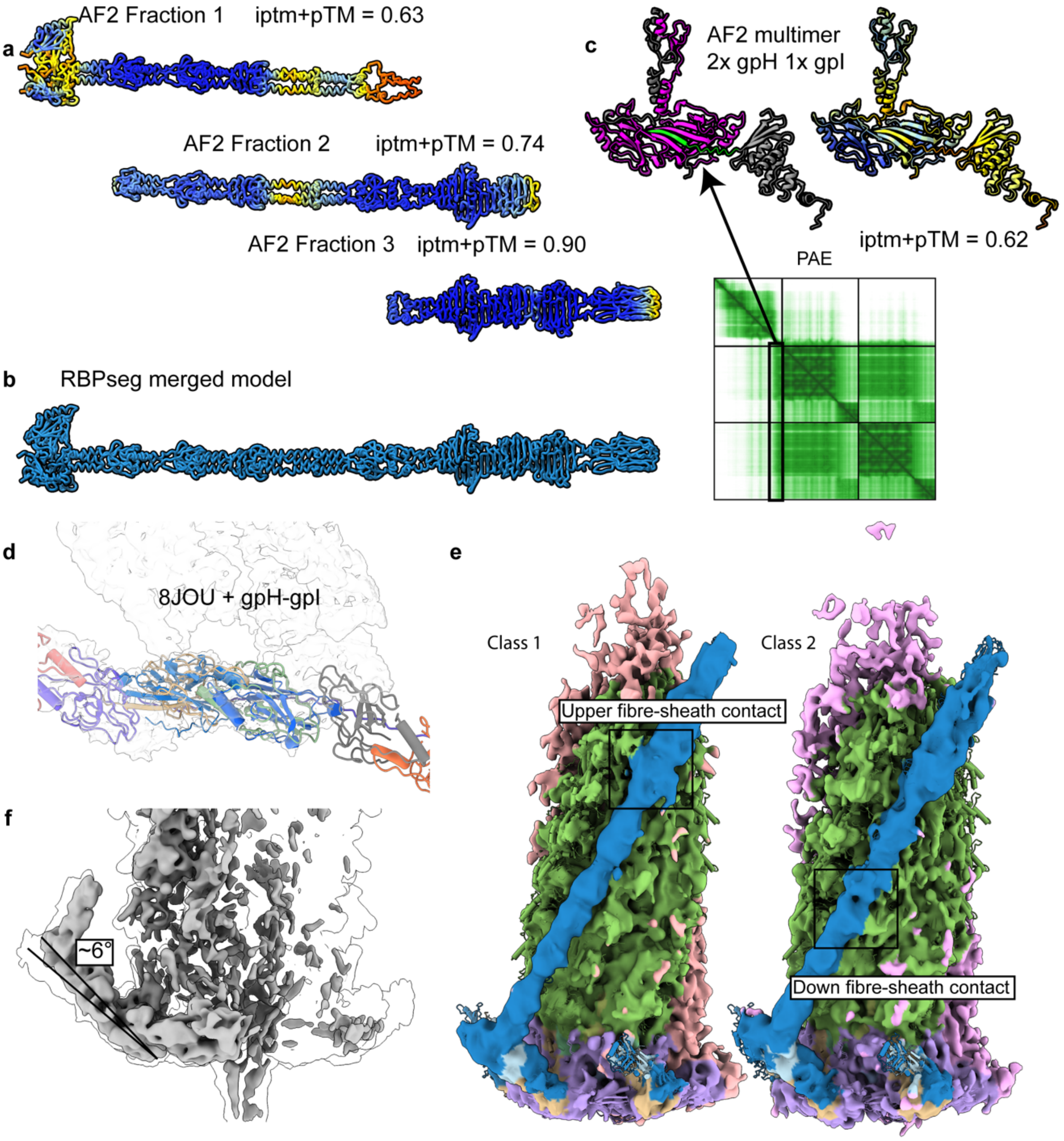


**Supplementary Figure 8. Fibre Alphafold confidence metrics and rope superposition.** (a) AlphaFold-predicted structural fractions of the fibre, coloured by per-residue confidence (pLDDT). (b) Built fibre model obtained by merging individual fractions using the RBPseg pipeline. (c) AlphaFold prediction of the gpH N-terminal region in complex with the triplex component gpI. On right: coloured by per-residue confidence (pLDDT). On left: colored based on the Predicted aligned error (PAE) plot region. (d) Rigid-body fitting of the rope model (PDB ID 8JOU) into the low-resolution P2 cryo-EM map with superposition of the proposed gpH-gpI model. (e) Two reconstructed classes for the upright conformation of the tail fibre, with squares showing contact points. (f) Superimposition of two classes map, showing the approximately 6° angle shift.

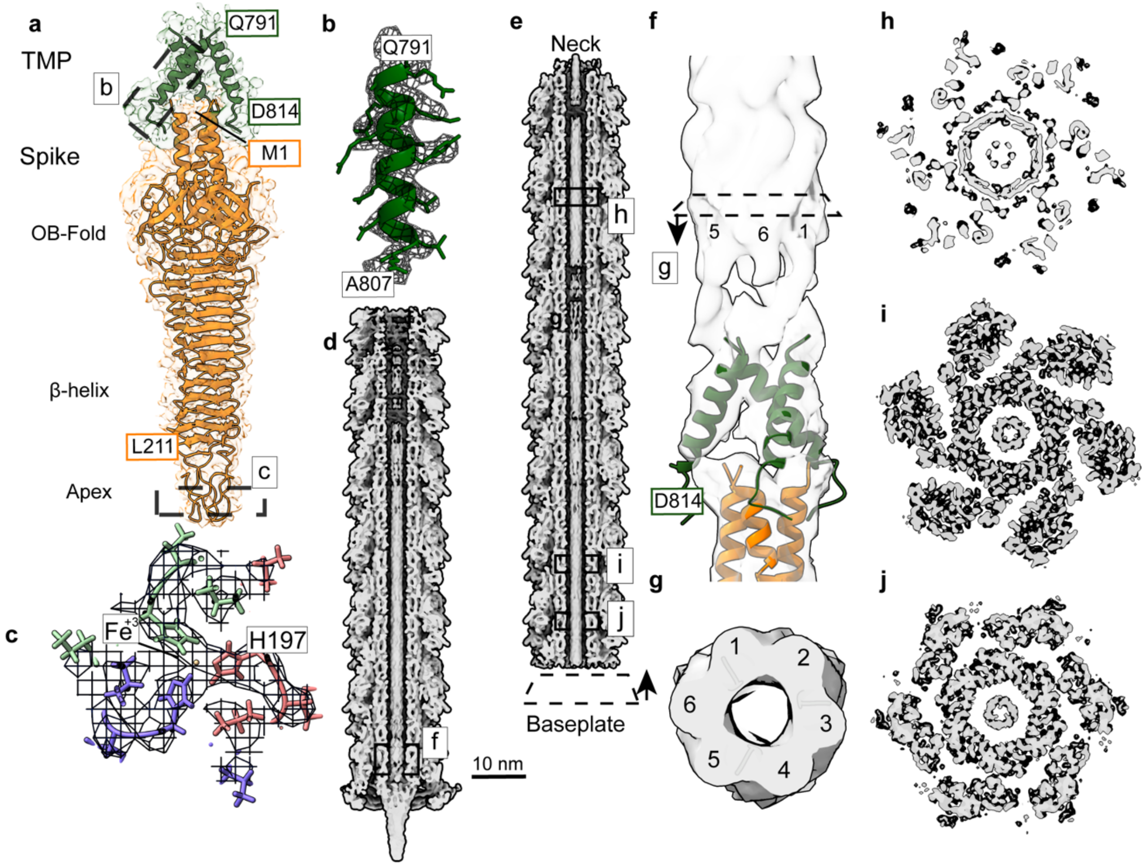


**Supplementary Figure 9. Structural analysis of the P2 spike and TMP C-terminus.**

(a) C3-refined local map of the P2 spike protein and the C-terminus of the TMP (29SG / EMD-57341) highlighting the major structural domains of the P2 spike. (b) Focus visualization of the Apex domain, with a locally sharpened map revealing internal density corresponding to an iron coordination site. (c) ModelAngelo prediction of the TMP C-terminus fitted into the sharpened helical density. (d-e) Longitudinal cross-section of P2 truck C3-local refined map of the TMP after low pass filtering. f) Focus view of (d) showing the density continuity between the TMP terminal and the map. (g) Horizontal cross-sections of the map in (f). (h-j) Horizontal cross-sections of the map in (e) at different at different hights.

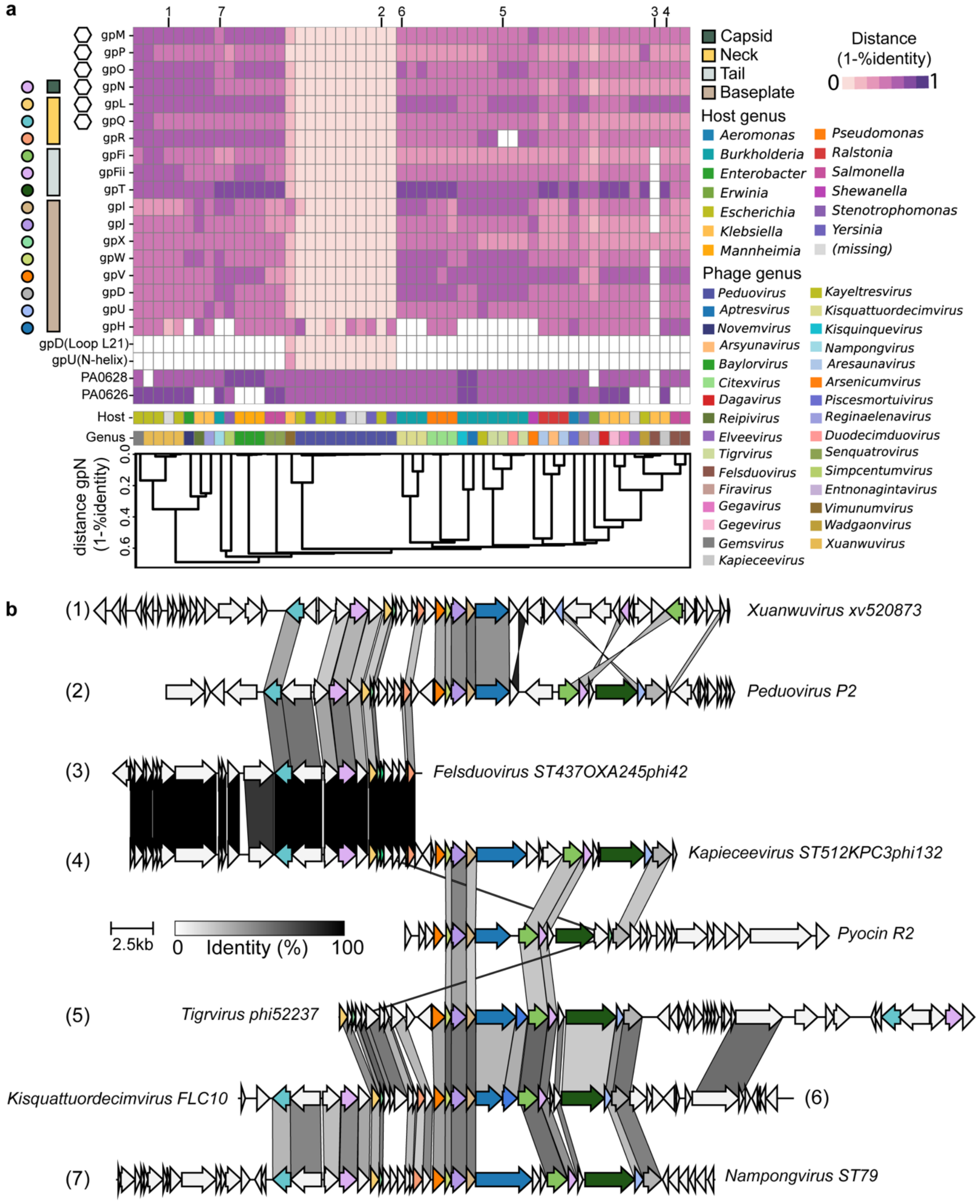


**Supplementary Figure 10. Sequence comparisons of phage P2 with other *Peduoviridae* phages.**

(a) Heatmap showing pairwise sequence similarity of conserved structural proteins and orthogroups between bacteriophage P2 and representative *Peduoviridae* phages. Hierarchical clustering was performed using pairwise sequence distances of the major capsid protein (gpN).

Phage host and genus is coloured as shown in legend. Proteins belonging to capsid, neck, tail truck and baseplate are also shown by color. Hexagonal markers represent P2 orthogroups. (b) Genome-scale alignment of selected phages highlighted in a, illustrating conservation of gene order and structural modules. Structural protein-encoding genes are color-coded according to orthogroup assignment. Grey links connect homologous proteins sharing at least 30% amino-acid sequence identity.

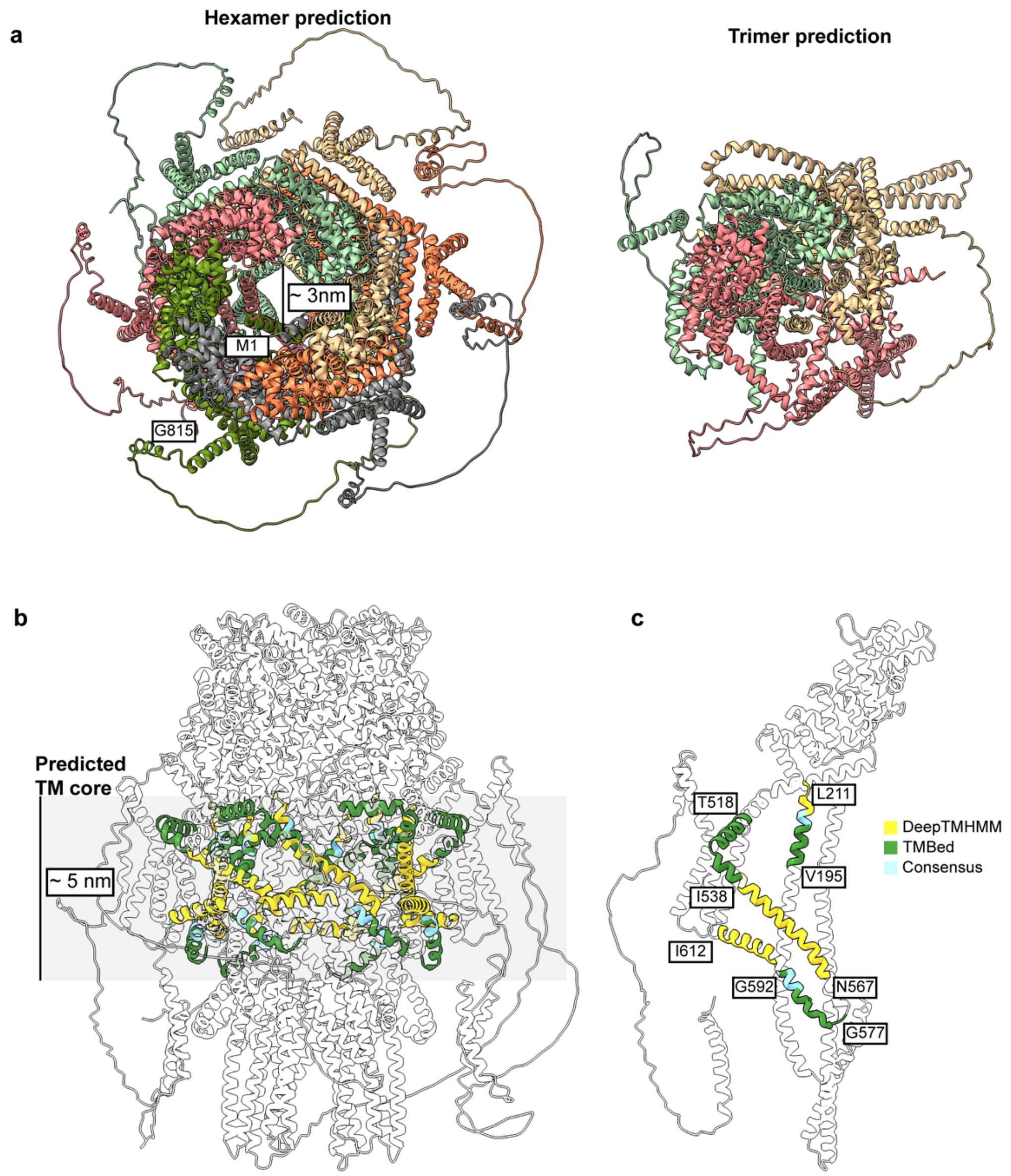


**Supplementary Figure 11. TMP AlphaFold predictions.**

(a) Hexamer AlphaFold2 prediction of P2's TMP (left, iptm+ptm: 0.69) and trimer AlphaFold2 predictions of P2's TMP (right, iptm+ptm: 0.48). (b) side view of hexamer prediction, highlight the predicted putative transmembrane helices. (c) Single chain representation of the hexamer prediction, highlighting transmembrane helices and their N and C-terminal residues.

**Supplementary Table 1: Cryo-EM data scollection**

| | Pre-contracted | | | | | | | | | Post-contraction | |
|---|---|---|---|---|---|---|---|---|---|---|---|
| | Capsid | Portal | Baseplate core | Baseplate wedge | Spike | Tail/TMP | Fibre two conformati ons | Fibre higher resolution | Fibre adaptor | Tail | Portal |
| **EMDB** | EMDB-57258 | EMDB-57255 | EMD-57254 | EMD-57254 | EMD-57341 | | EMD-57256 | EMD-57256 | EMD-57256 | EMD-57343 | EMD-57344 |
| **PDB** | 29NY | 29NE | 29ND | 29ND | 29SG | | - | - | - | - | - |
| **EMPIAR** | | | | | | | | | | | |
| Magnification | 105,000 | 105,000 | 105,000 | 105,000 | 105,000 | 105,000 | 105,000 | 105,000 | 105,000 | 105,000 | 105,000 |
| Voltage (kV) | 300 | 300 | 300 | 300 | 300 | 300 | 300 | 300 | 300 | 300 | 300 |
| Electron exposure (e–/Å$^2$) | 50 | 50 | 50 | 50 | 50 | 50 | 50 | 42 | 42 | 42 | 42 |
| Defocus range (µm) | [-2.5;-0.5] | [-2.5;-0.5] | [-2.5;-0.5] | [-2.5;-0.5] | [-2.5;-0.5] | [-2.5;-0.5] | [-2.5;-0.5] | [-2.5;-0.5] | [-2.5;-0.5] | [-2.5;-0.5] | [-2.5;-0.5] |
| Pixel size (Å) | 1.2 | 1.2 | 1.2 | 1.2 | 1.2 | 1.2 | 1.2 | 1.2 | 1.2 | 1.2 | 1.2 |
| Symmetry imposed | I/C1 | C6 | C3 | C1 | C1 | C3 | C1 | C1/C3 | C1 | C6 | C6 |
| Initial particle images (no.) | 8,598 | 37,941 | 51,583 | 38,184 | 51,583 | 38,184 | 38,320 | 38,320 | 40,892 | 18,179 | - |
| Final particle images (no.) | 7,986 | 14,634 | 38,184 | 51,942 (symmetry expanded) | 153,716 (symmetry expanded) | 18,886 (down tail) 6,186 (upper tail) | 19,176 (class 1) 19,144 (class 2) | 38,320 | 27,231 | 17,591 | - |
| Map resolution (Å) (FSC threshold = 0.143) - mask | 09.46 | 2.73 | 2.83 | 3 | 2.94 | 2.88 (down) 4.57 (upper) | 6.00 | 5.42 (C1) 5.67 (C3) | 4.85 | 09.46 | 09.46 |
